# Reaction-time asymmetry covaries with shifts in hand choice in macaques

**DOI:** 10.64898/2026.10.27.762253

**Authors:** Charles D. Holmes, Suhwan Hong, Sergei Gepshtein, Lawrence H. Snyder, Eric Mooshagian

## Abstract

How primates choose which hand to reach with remains poorly understood. Humans prefer the hand on the target’s side of the body midline, and that preference shifts with biomechanical cost and with head and gaze direction. Earlier monkey studies used only two targets, one per side, too few to fit a psychometric function. Reaction time may also reflect the choice; in humans it is longest at the point of subjective equality (PSE), the target position at which either hand is equally likely to be chosen. We measured hand choice at ten horizontal target positions in two male rhesus macaques (*Macaca mulatta*) that chose freely on each trial. For peripheral targets, both animals used the hand on the same side of the body midline, with a well-defined transition in between. Displacing the hands, rotating the head, or displacing gaze to one side each moved the PSE toward that side. Reaction time was longest near the PSE. Hand displacement slowed the hand on the displaced side, and gaze displacement slowed the hand opposite the gaze. In every condition, the farther the PSE lay toward one side, the slower that side’s hand was relative to the other. Macaque hand choice thus resembles human hand choice, making the macaque a tractable model for studying how the brain chooses a hand.

**New & Noteworthy:** Macaques choosing freely between hands mostly use the hand on the target’s side. The point of subjective equality, where either hand is equally likely, moves with hand displacement, head rotation, and gaze displacement, and reaction time is longest there. In every condition, the farther this point lies toward one hand’s side, the slower that hand is relative to the other. The macaque is thus a tractable model for studying how the brain chooses a hand.

## Introduction

Choosing which hand to use is a frequent everyday decision. Many reaches can be made with either hand, and for these the brain must choose one. Target location, task demands, biomechanical cost, and individual hand preference all influence which hand is chosen (Gabbard and Rabb 2000; Helbig and Gabbard 2004; Hirayama et al. 2022). Humans tend to reach to targets on the left with the left hand and to targets on the right with the right hand, with a transition zone near the body’s midline across which the chosen hand switches (Bryden et al. 2000; Stoloff et al. 2011). The center of that zone, the target position at which either hand is equally likely to be chosen, is the point of subjective equality (PSE). The effort a reach requires changes how often each hand is chosen (Kim et al. 2022; Wang et al. 2021) and thus moves the PSE (Bakker et al. 2017; McDermott and Himmelbach 2019). Displacing the hands changes how far each hand must reach to a given target, and so changes the effort of each reach. Rotating the head or redirecting gaze changes neither the effort of a reach nor the target’s position relative to the body, yet it still moves the PSE (Bakker et al. 2018). Rotating the head moves the target in head-centered coordinates, and redirecting gaze moves it in eye-centered coordinates, so hand choice appears to draw on more than one reference frame (Bakker et al. 2018).

Reaction time may also carry information about the choice. In human reaching, responses are slowest to targets near the PSE (Oliveira et al. 2010) or the midline (Stoloff et al. 2011). Lowering the effort a reach requires or raising its reward increases both how often it is chosen and how quickly the movement begins in humans (Wang et al. 2021). In monkeys, saccades to the target chosen more often also start sooner (Milstein and Dorris 2011). The one test in monkey reaching found no such relation in reaction time, but it measured reaction time on trials where the hand was instructed (Dancause and Schieber 2010). If the saccade result holds for the hands, the hand chosen more often should start its reach sooner, and the difference between the two hands’ reaction times should vary with the PSE. Human reaching shows such a between-hand asymmetry in reaction time under free choice but not when the hand is assigned (Dexheimer et al. 2022).

How the brain combines these influences to select one hand is unknown. Macaques, in which single neurons can be recorded and brain areas perturbed reversibly, are the principal model for the neural control of reaching, but that work has concentrated on instructed movements (Churchland et al. 2012; Georgopoulos et al. 1986; Mooshagian et al. 2021). Hand preference in primates has been surveyed extensively (Papademetriou et al. 2005), but few studies have examined how it varies with target location, and those presented the target at one of two locations, one on each side of the midline (Dancause and Schieber 2010; Gardinier et al. 2006; Lee and Schieber 2006). In those studies, hand choice depended more on each monkey’s preferred hand than on the target’s side, and the animals often reached over the midline to use it (Dancause and Schieber 2010; Gardinier et al. 2006). When head and gaze direction were varied instead, with target position held fixed, the monkeys chose the hand and target ipsilateral to head direction (Dancause and Schieber 2010). Whether macaques, like humans, primarily choose the hand on the target’s side, with a well-defined transition between the hands, was therefore unclear. Because hand choice depends on where the target lies, neural activity that varies with target position could reflect the target, the chosen hand, or both. Knowing how the use of each hand varies across target positions, and how displacing the hands or gaze or rotating the head shifts that pattern, would help separate activity related to target location from activity related to hand choice.

Here we asked three questions in macaques that freely chose which hand to use in a visually guided reaching task adapted from human psychophysics. First, if macaques choose a hand as humans do, they should use the hand on the target’s side, with a transition zone between the hands. Second, if hand choice depends on more than the target’s position relative to the body, displacing the hands, rotating the head, or displacing gaze should each move the PSE; if it depends on that position alone, head rotation and gaze displacement would leave the PSE in place. Third, if hand choice and reaction time depend on a common process, reaction time should be longest near the PSE, where the choice is most uncertain. We also tested, post hoc, whether the RT asymmetry, our term for the percentage by which right-hand reaction time exceeds left-hand reaction time in the same session, varies with the PSE, as predicted above, even when the hands, head, and gaze are held in place. For peripheral targets, both macaques used the hand on the same side of the body midline, with a well-defined transition between the hands, and each manipulation moved the PSE in the direction of the manipulation. Reaction time was longest for targets near the PSE. Hand and gaze displacement also changed reaction time; rightward gaze displacement, which moved the PSE rightward, lowered the RT asymmetry. In a post hoc analysis, the RT asymmetry rose as the PSE moved rightward, and the slope was positive within every condition.

## Results

Two rhesus macaques (*Macaca mulatta*) were trained to perform a visually guided reaching task in which they freely chose which hand to use (Fig. 1A, B). At trial onset, the animal rested its left and right hands on separate home buttons, 9 cm to either side of midline (the body’s midsagittal plane), with its head fixed facing forward, and fixated a central point on a touchscreen. After a 300 ms hold, a target appeared at one of 30 locations, ten horizontal positions from −27 to +27 deg from the midline across both hemifields crossed with three vertical rows at −18, −4 and +10 deg relative to the fixation point (Fig. 1A; Methods). The animal reached for and touched the target (reaching accuracy, Supplemental Figure S1). Either hand could be used, but the other hand had to stay on its home button, so that the animal used exactly one hand on each trial.

**Figure 1:**
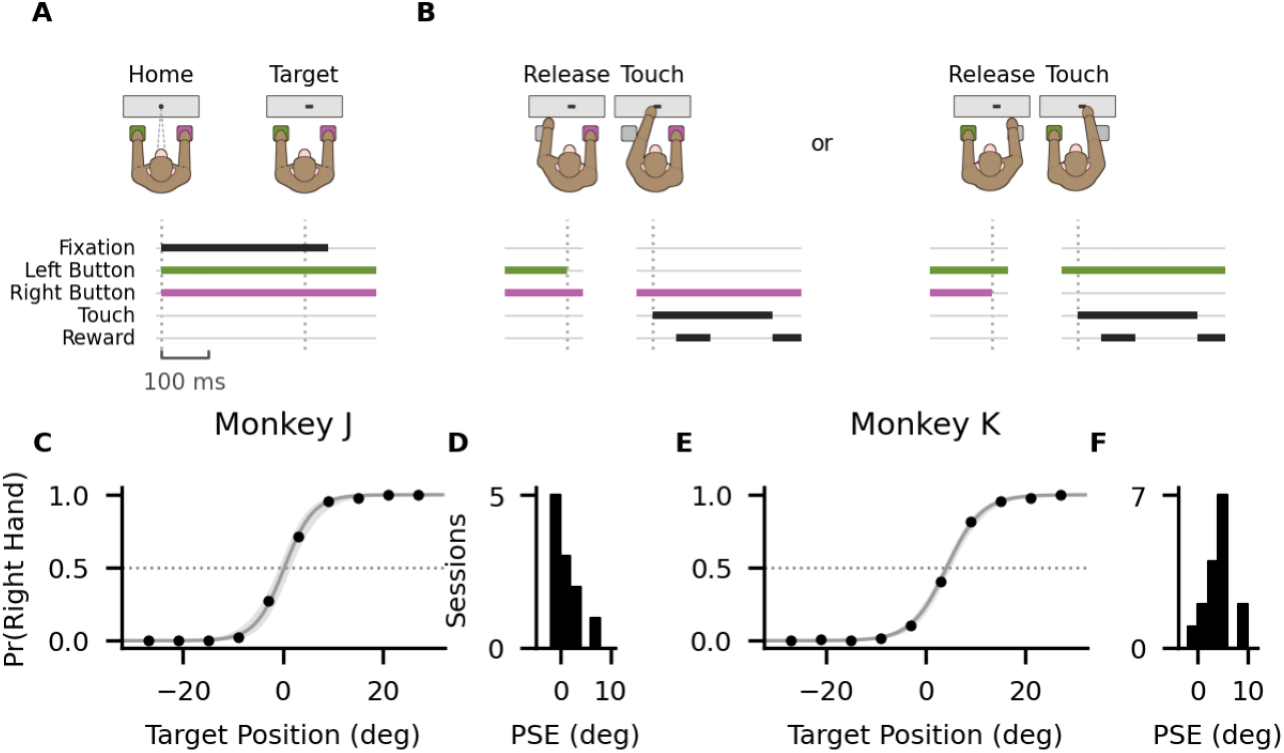
Hand choice depends on target position. (A, B) The free-choice reaching task, with the start of a trial (A) and its two possible endings (B), a reach with the left hand or with the right hand. Vertical dotted lines mark the time of the event drawn above each. (A) The animal placed each hand on its own home button and fixated a central fixation point (Home). After 300 ms, a target appeared at one of 30 locations (Target; Methods); fixation was enforced for 50 ms more, after which eye position was unconstrained. (B) The animal lifted one hand, and only one, from its home button (Release) and touched the target with it (Touch). Ribbons show the behavior: black, the required fixation (Fixation), the hand on the target (Touch), and the water reward (Reward), small after the touch and larger once the touch had been held; green and purple, the left and right hand on its home button, respectively (Left Button, Right Button). In the drawings, a held button is green or purple and a released one gray. The break after A omits part of the reaction time (Target to Release), and the gaps within B omit part of the movement time (Release to Touch); both varied across trials. Scale bar, 100 ms. (C, E) Proportion of right-hand reaches against horizontal target position in one example session per animal. Points, observed proportions; gray line, logistic fit; shading, its 95% confidence interval; horizontal dotted line, a proportion of 0.5. The point of subjective equality (PSE) is where the fit crosses it. (D, F) Distribution of session PSEs without a manipulation, one per session fitted to all of its trials, for Monkey J (11 sessions) and Monkey K (16 sessions).

Monkey J performed the task in 37 sessions and Monkey K in 35, one per day, each made up of blocks of 150–300 consecutive trials. On 11 of Monkey J’s sessions and 16 of Monkey K’s, the home buttons, head, and fixation point stayed as described above in every block; these were the sessions without a manipulation. On each of the rest, we manipulated one of the three across blocks, displacing the hands (11 sessions in Monkey J, 8 in Monkey K), rotating the head (10 and 5), or displacing gaze (5 and 7); on one Monkey K session we manipulated both the hands and the head (Methods).

### Hand choice depends on target position

Both animals chose to use the left hand for targets on the left and the right hand for targets on the right, with a transition zone between the hands near the midline (Fig. 1C, E). On the sessions without a manipulation, Monkey J completed 28 blocks and Monkey K completed 49 blocks. Every block of a session had the same configuration, so we fit one logistic function per session, to all of its trials, to estimate the PSE (example sessions, Fig. 1C, E). Across sessions, Monkey J’s median PSE was 0.29 deg (95% CI [-0.88, 3.0]), and Monkey K’s was 4.2 deg (95% CI [3.1, 4.9]), a difference of 3.9 deg (95% CI [1.1, 5.3]). The 10–90% transition width, the span of target positions over which the probability of a right-hand reach rose from 10% to 90%, was 11 deg in Monkey J (95% CI [9.5, 13]) and 13 deg in Monkey K (95% CI [11, 14]), a difference of 1.9 deg (Monkey K minus Monkey J; 95% CI [-1.7, 3.6]). The animals therefore differed in the PSE, and the interval for the width difference included zero. Both medians lay right of the midline, but the 95% CI excluded the midline only in Monkey K.

Both animals had prior training on instructed bimanual reaches, so their use of either hand could be a product of that history rather than a general tendency. We therefore tested a third animal, Monkey A, who lacked that history. Monkey A had been trained to reach to a touch screen, but with only one assigned hand per session, a history that predicts persistent use of one hand. Monkey A was also naive to home buttons, so we used a modified task (Methods). In every session (5 of 5), the proportion of right-hand reaches rose the farther right the target was (median PSE 8.6 deg, 95% CI [7.1, 20]; Supplemental Figure S2). Split use of both hands across target positions therefore does not require a history of bimanual training.

### Displacing the hands or gaze or rotating the head shifts hand choice

To test whether hand choice depends only on the target’s position relative to the body, we displaced the hands, rotated the head, or displaced gaze (Fig. 2A–C). We kept the home-button pair centered or displaced it laterally by 3.5 or 4.5 cm to either side, rotated the head to −15, 0, or +15 deg with body and hands fixed, and moved the fixation point to −15, 0, or +15 deg with head and body centered (reaching accuracy, Supplemental Figures S3–S8). Each session with a manipulation tested at least two of these shift levels.

**Figure 2:**
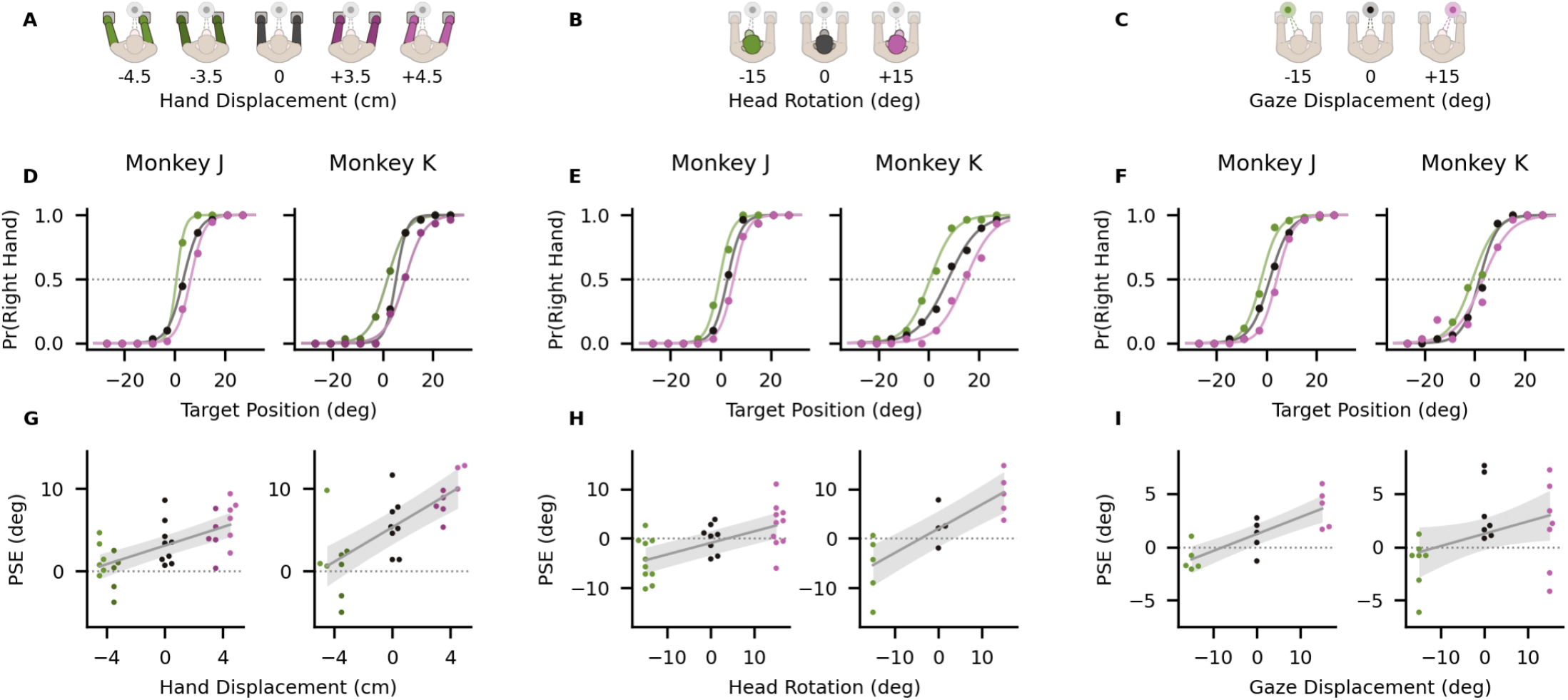
The PSE moves in the direction of each manipulation. Hand displacement (A, D, G), head rotation (B, E, H), and gaze displacement (C, F, I), one per column. (A–C) Schematics, seen from above, with the fixation point drawn nearer than to scale. The part each manipulation moves is filled in the color of its shift (black, no shift); the rest of the animal is faded, and so is the fixation point where gaze was not moved. The home-button pair was displaced laterally, by ±3.5 or ±4.5 cm; the head was rotated to −15, 0, or +15 deg with body and hands fixed; and gaze was displaced to −15, 0, or +15 deg with head and body fixed. Each drawing is labeled with its shift, and its color marks that shift throughout its column. (D–F) Proportion of right-hand reaches against horizontal target position in one example session per animal. Points, observed proportions; curves, one logistic fit per shift level, to all of that level’s trials in the session. Each fit crosses the dotted line, a proportion of 0.5, at its point of subjective equality (PSE). (G–I) The PSE at each shift level of each session against the shift, Monkey J left and Monkey K right. Points, one per session and shift level (hand displacement, 31 shift levels over 11 sessions in Monkey J and 24 over 8 in Monkey K; head rotation, 28 over 10 and 14 over 5; gaze displacement, 15 over 5 and 21 over 7). Points sharing a shift are offset horizontally so that none covers another; the offset carries no information. Gray line, linear fit with a random intercept per session (Methods); shading, its 95% confidence interval; dotted line, zero PSE.

Within single sessions (Fig. 2D–F) and across sessions (Fig. 2G–I), the PSE moved in the direction of each manipulation. Hand displacement moved the PSE +0.56 deg per cm in Monkey J (95% CI [0.38, 0.75]; n = 31 shift levels over 11 sessions) and +1.1 deg per cm in Monkey K ([0.76, 1.3]; n = 24; Fig. 2G). Head rotation moved it +0.23 deg per deg in Monkey J ([0.18, 0.29]; n = 28) and +0.49 deg per deg in Monkey K ([0.35, 0.63]; n = 14; Fig. 2H), and gaze displacement moved it +0.16 deg per deg in Monkey J ([0.11, 0.21]; n = 15) and +0.12 deg per deg in Monkey K ([-0.0052, 0.24]; n = 21; Fig. 2I). Every slope was positive, and all but Monkey K’s gaze slope were significantly greater than zero (two-sided Wald test; p < 0.001 in each case except Monkey K’s gaze slope, p = 0.06), and the four head and gaze slopes were all significantly less than one, the value expected if the PSE moved by as many degrees as the head was rotated or gaze was displaced (p < 0.001 for each). Monkey K’s gaze slope did not differ from Monkey J’s (difference -0.043 deg per deg, 95% CI [-0.19, 0.11]; p = 0.57), and pooled across the two animals the gaze slope was +0.13 deg per deg (95% CI [0.060, 0.21]; p < 0.001). Hand choice therefore did not depend only on the target’s position relative to the body. The PSE also depended on the position of the hands and on the orientation of the head and eyes.

### Reaction time is longest near the PSE

Reaction time (RT), the time from target onset until the reaching hand left its home button, was longest for targets near the PSE (Supplemental Figure S9A, B). In most sessions, reaches were slower to near-PSE targets (within 6 deg of that session’s PSE) than to targets at least 15 deg to that hand’s side of the PSE, in both animals and for both hands (Monkey J, all 11 sessions for each hand; Monkey K, 14 of 16 for the left hand and 13 of 16 for the right; Fig. 3A, B). Paired sign-flip permutation tests over sessions, one per animal and hand, put this near-PSE RT increase at 6.9% (95% CI [4.6, 9.2]) and 7.6% ([4.6, 10.6]) in Monkey J (left and right hand, n = 11 sessions; both p < 0.001) and at 9.5% ([4.6, 14.7]) and 3.8% ([0.7, 7.0]) in Monkey K (n = 16 sessions; p < 0.001 and p = 0.004; p values uncorrected, corrected in Supplemental Table S3). Pooled across the two animals, the increase was significant for both hands under every manipulation (Fig. 3F, I, L; paired sign-flip tests over sessions, Benjamini–Hochberg corrected; largest p = 0.004), and the animals did not differ after correction (smallest p = 0.15), although Monkey K’s increase was smaller than Monkey J’s in seven of the eight combinations of hand and condition, most clearly under gaze displacement. Tested in each animal alone, the mean increase was positive in 12 of the 12 combinations of animal, hand, and manipulation and significant in 5 (Supplemental Table S3). In 3 of the non-significant combinations every session showed an increase, but each had only five sessions, too few for the test to reach significance; the other four were all Monkey K’s.

**Figure 3:**
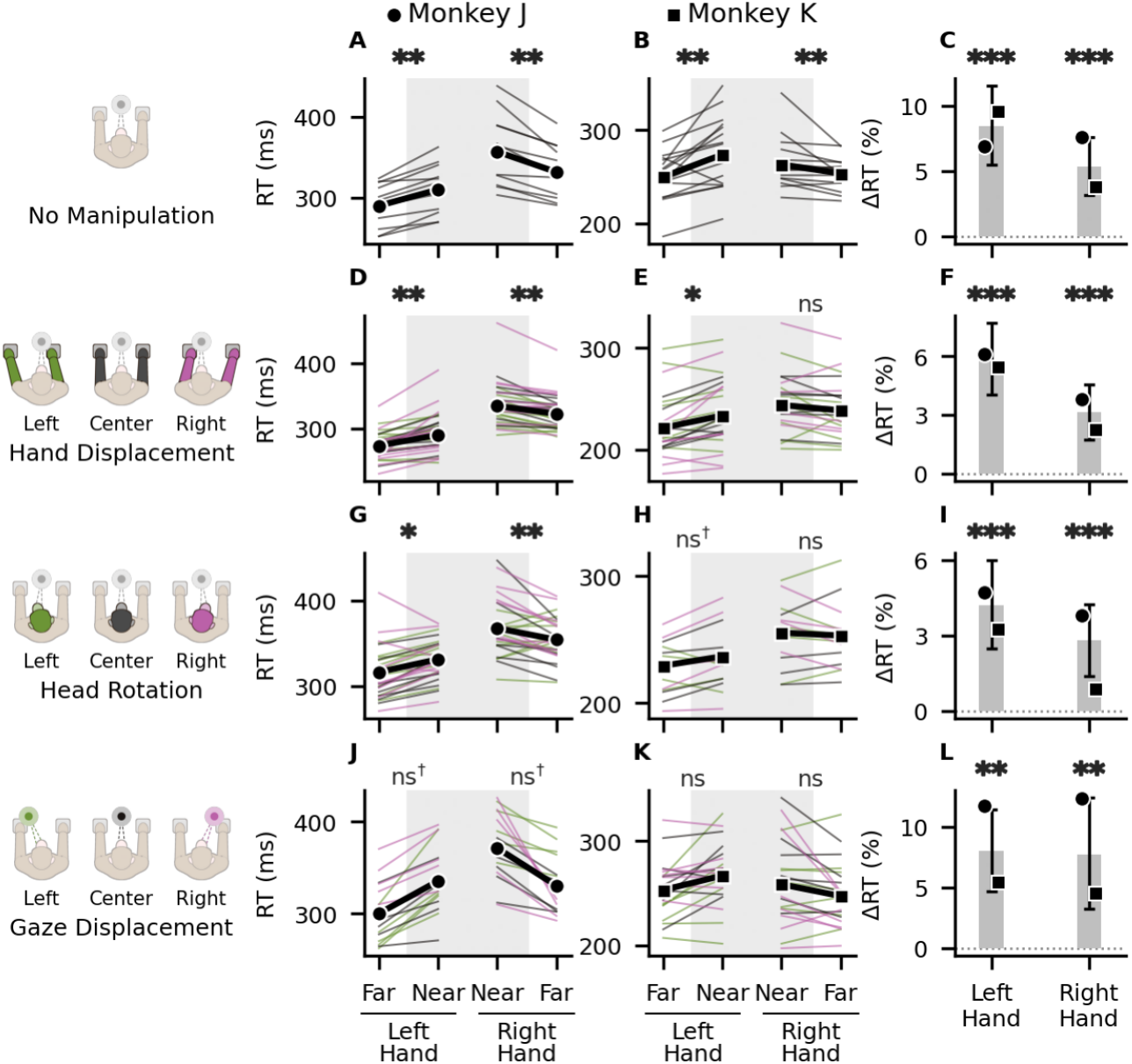
Reaction time is longest near the PSE. Reaction time (RT) near the point of subjective equality (PSE) and far from it, with no manipulation (A–C), hand displacement (D–F), head rotation (G–I), and gaze displacement (J–L), one per row. At the left of each row, an icon for each shift direction, drawn as in Fig. 2 and labeled with that direction. (A, B, D, E, G, H, J, K) Each hand’s geometric-mean RT to targets within 6 deg of the PSE of that session and shift level (Near, shaded) and to targets at least 15 deg to that hand’s side of it (Far), for Monkey J (left column) and Monkey K (middle column). Thin lines, one per session and shift level, colored by the direction of the shift (green, leftward; black, centered or none; purple, rightward); the hand-displacement row pools both displacement sizes in each direction. Lines with no far target are omitted (Methods). Thick black line, the mean over sessions, each session’s shift levels averaged first (circle, Monkey J; square, Monkey K). Marks over each hand’s pair, the significance of that hand’s near-PSE RT increase in that animal, the percentage by which its Near RT exceeds its Far RT (paired sign-flip test over sessions), Benjamini–Hochberg corrected across the 16 per-animal tests (*, p < 0.05; **, p < 0.01; ns, not significant; ns†, every session’s increase was positive, but five sessions cannot reach p < 0.05). (C, F, I, L) The near-PSE RT increase (ΔRT). Bars, the mean pooled across the two animals, every session weighted equally, with its 95% confidence interval (t distribution, uncorrected); circle and square, each animal’s mean. Dotted line, no increase. Marks, the significance of each pooled increase (paired sign-flip test over sessions), Benjamini–Hochberg corrected across the eight bars (**, p < 0.01; ***, p < 0.001). The sessions without a manipulation are those of Fig. 1D, F, and the sessions and shift levels under one those of Fig. 2G–I. Session counts and per-animal tests are in Supplemental Table S3.

### Reaction time under each manipulation

The manipulations also changed reaction time (Fig. 4). Each effect was measured one hand at a time, on that hand’s reaches to peripheral targets on its own side (at least 15 deg from the body midline). Reaches to these targets were made with the hand on the target’s side on 99.3% of trials in the no-manipulation sessions, and on 98.5, 97.8 and 98.8% under the hand, head and gaze manipulations, so hand choice at these targets changed little across conditions. Each effect was tested with a paired sign-flip permutation test over sessions, and its p value is corrected for multiple comparisons (Methods).

**Figure 4:**
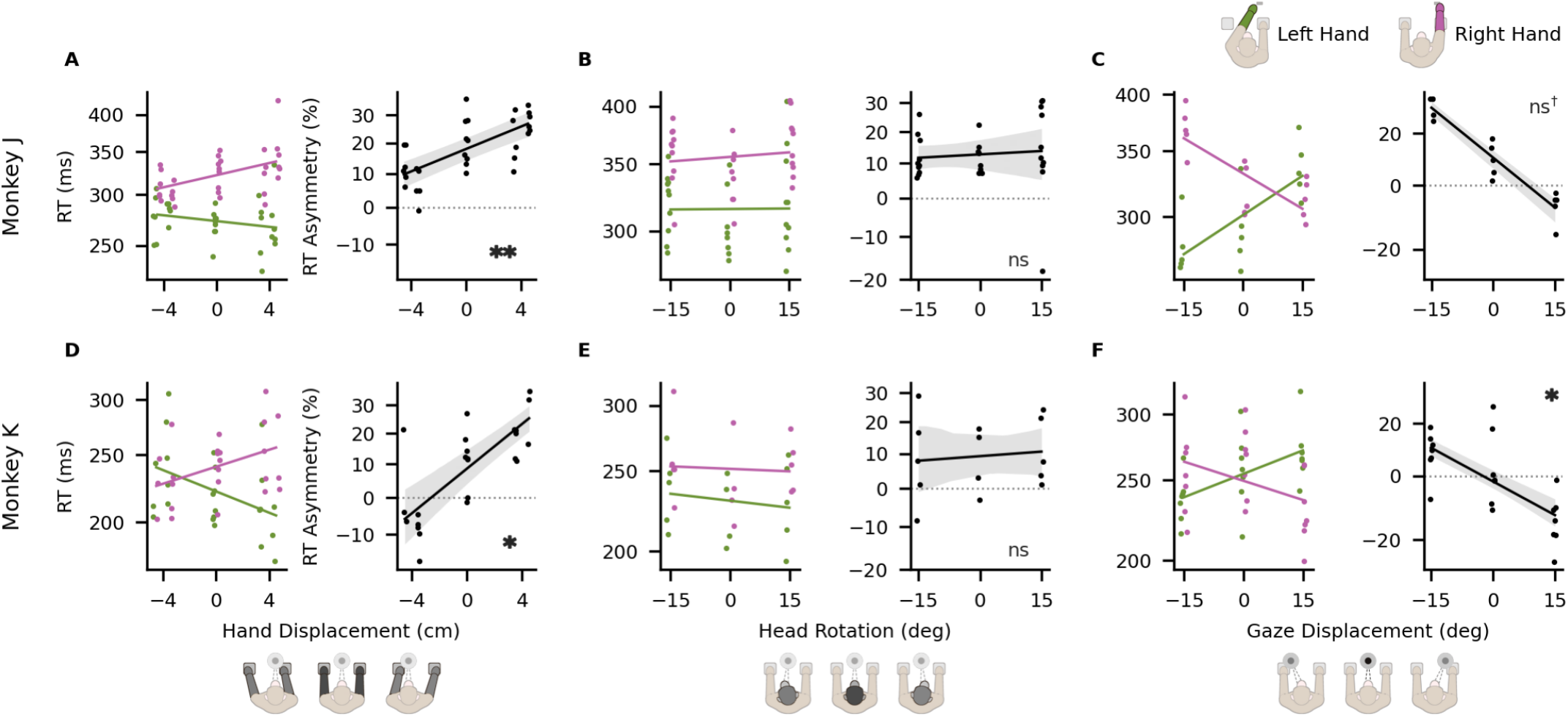
Hand and gaze displacement change the RT asymmetry. Hand displacement (A, D), head rotation (B, E), and gaze displacement (C, F), one per column, for Monkey J (A–C) and Monkey K (D–F). Icons under each column as in Fig. 2. Left panel of each lettered pair, reaction time (RT) of the left hand (green) and the right hand (purple) at peripheral targets on that hand’s own side (at least 15 deg from the body midline), the geometric mean at each shift level of each session, on a logarithmic axis labeled in ms. Points, one per hand, session and shift level, offset horizontally for visibility, each hand toward its own side; lines, each hand’s least-squares slope within each session averaged across sessions. Right panel, the RT asymmetry, the percentage by which right-hand RT exceeds left-hand RT at the same shift level of the same session. Black line, its slope, fitted the same way; shading, its 95% confidence band (Methods); dotted line, zero asymmetry. Marks in each asymmetry panel, the significance of that slope (paired sign-flip test over sessions), Benjamini–Hochberg corrected across the six asymmetry panels (*, p < 0.05; **, p < 0.01; ns, not significant; ns†, every session’s slope had the same sign, but five sessions cannot reach p < 0.05). Sessions and shift levels are those of Fig. 2G–I.

Displacing the hands toward one side lengthened the reaction time of the hand on that side in all four hands, though no hand’s trend was significant in either animal alone after correction (Fig. 4A, D). Reaction time rose by 1.1% per cm for Monkey J’s right hand (p = 0.07), by 1.8% per cm for Monkey K’s left hand (p = 0.07), by 1.4% per cm for Monkey K’s right hand (p = 0.07), and by 0.52% per cm for Monkey J’s left hand (p = 0.35). Pooled across the two animals, both hands’ trends were significant (p = 0.003 for the left hand and p = 0.001 for the right), and the animals did not differ (smallest p = 0.18). The RT asymmetry, the percentage by which right-hand reaction time exceeds left-hand reaction time in the same session and shift level, rose with rightward displacement by 1.6% per cm in Monkey J and 3.2% per cm in Monkey K (Fig. 4A, D), with the same sign in every session of both animals (p = 0.006 and p = 0.02).

Rotating the head in either direction slowed both of Monkey J’s hands, though not significantly after correction (Fig. 4B, E): the right hand was 6.8% slower with the head rotated than with it centered (p = 0.07), and the left hand 3.1% slower (p = 0.26). Monkey K’s hands slowed by 2.3% (right) and 2.3% (left), but only 4 of its sessions had a centered block, too few for any effect to reach significance (smallest attainable p = 0.12). Neither hand’s reaction time depended on the direction of rotation in Monkey J (p = 0.95 for the left hand and p = 0.75 for the right) or pooled across the two animals (p = 0.69 for the left hand and p = 0.69 for the right), and neither did the RT asymmetry, which changed by 0.062% per deg in Monkey J and 0.083% in Monkey K (p = 0.75 and p = 0.75).

Displacing gaze had the opposite effect to displacing the hands, lengthening each hand’s reaction time as gaze moved away from that hand’s side (Fig. 4C, F). In Monkey J, reaction time rose by 0.61% per deg for the left hand and by 0.55% per deg for the right, with every session showing that sign (p = 0.17 for each; no test on 5 sessions can give less than 0.06, even before correction). In Monkey K, reaction time rose by 0.43% per deg for the left hand (p = 0.12) and by 0.35% per deg for the right hand (p = 0.07). Pooled across the two animals, both hands’ trends were significant (p = 0.002 for the left hand and p = 0.001 for the right), and the animals did not differ (smallest p = 0.49). The RT asymmetry therefore fell with rightward gaze displacement, by 1.2% per deg in Monkey J and 0.79% per deg in Monkey K (Fig. 4C, F), with the same sign in every session of both animals (p = 0.09 and p = 0.03).

Thus, across conditions, RT asymmetry did not consistently track the PSE, which every manipulation moved toward the side of the shift.

### RT asymmetry covaries with the PSE

That the RT asymmetry did not consistently track the PSE across conditions does not rule out a relation within a condition, where the PSE still varied from session to session. If hand choice and reaction time depend on a common process, sessions of the same condition whose PSE lay farther right should show a larger RT asymmetry.

Without a manipulation, the RT asymmetry was larger in sessions whose PSE lay farther right (Fig. 5A, B), rising by 1.9% per deg in Monkey J (95% CI [0.75, 3.0]; p = 0.005; n = 11 sessions) and by 3.0% per deg in Monkey K (95% CI [1.7, 4.3]; p < 0.001; n = 16 sessions). The analysis was post hoc, and with one PSE per session the slope was fitted by ordinary least squares (Methods). The RT asymmetry was measured at peripheral targets (at least 15 deg from the body midline), which carry the least information about the PSE (Methods).

**Figure 5:**
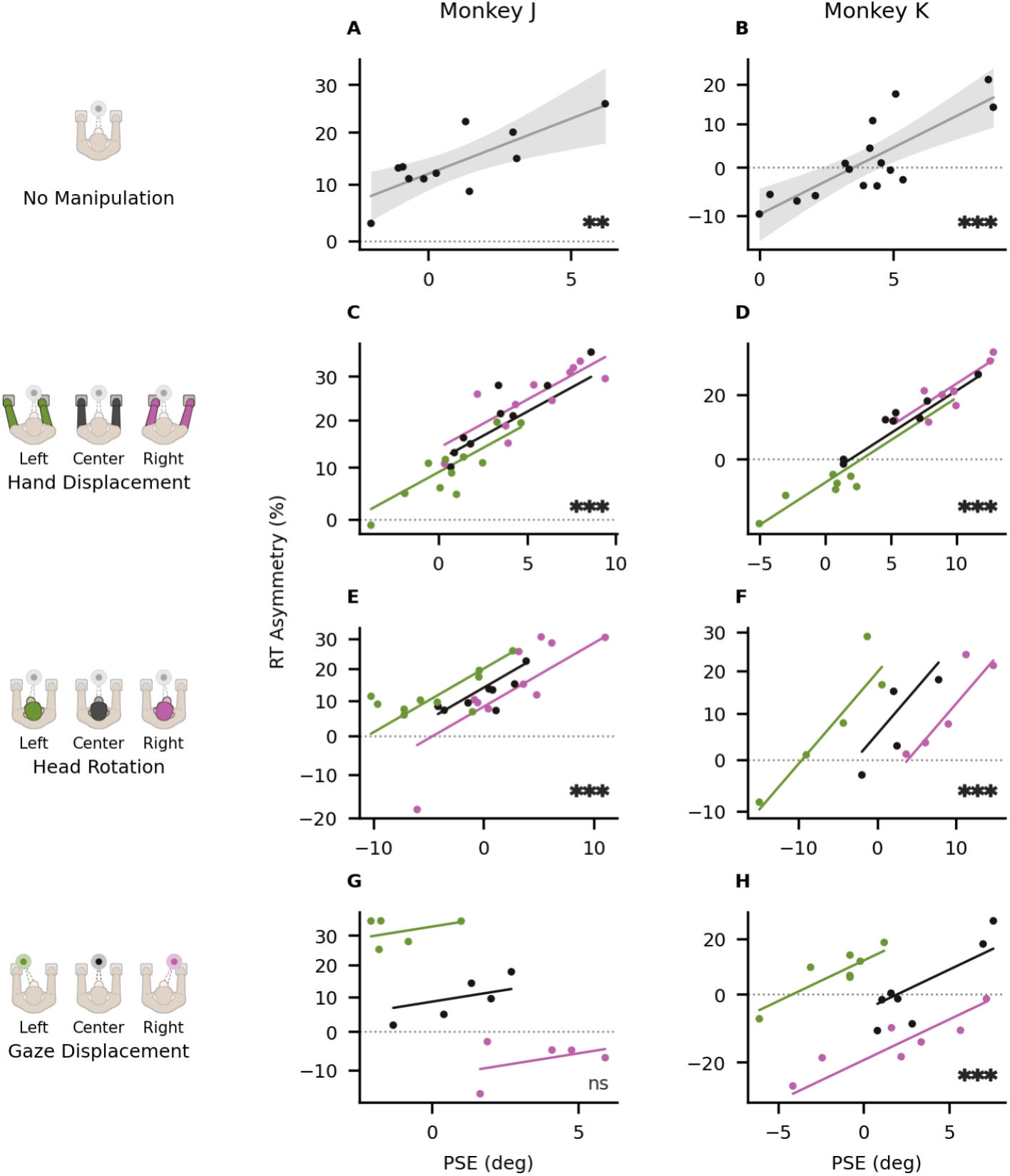
The RT asymmetry correlates with the PSE. Reaction time (RT) asymmetry, the percentage by which right-hand RT exceeds left-hand RT at targets on each hand’s own side at least 15 deg from the body midline, against the point of subjective equality (PSE), with no manipulation (A, B), hand displacement (C, D), head rotation (E, F), and gaze displacement (G, H), one per row, for Monkey J (left column) and Monkey K (right column). At the left of each row, an icon for each shift direction, drawn as in Fig. 2 and labeled with that direction. (A, B) Points, one per session, the sessions of Fig. 1D, F. Gray line, the least-squares linear fit across sessions; shading, its 95% confidence interval. (C–H) Points, one per session and shift level, the sessions and shift levels of Fig. 2G–I, colored by the direction of the shift (green, leftward; black, centered; purple, rightward); the hand-displacement row pools both displacement sizes in each direction. Lines, the fit across all of the manipulation’s sessions and shift levels, with the shift as a covariate and a random intercept per session; it has one slope, so it is drawn as parallel lines, one for each direction at that direction’s mean shift and across its PSE range. Marks, the significance of each panel’s slope, Benjamini–Hochberg corrected across the eight panels (**, p < 0.01; ***, p < 0.001; ns, not significant).

Under each manipulation, the RT asymmetry covaried with the PSE across sessions, with a positive slope in every condition in both animals (Fig. 5C–H; Supplemental Table S1). Hand and gaze displacement also changed the RT asymmetry across conditions (Fig. 4A, C, D, F), so the slopes that follow were fitted with the manipulation’s signed shift as a covariate and a random intercept per session (Methods). Supplemental Table S2 gives each fit without the covariate. For hand displacement and head rotation, the covariate changed neither the sign nor the significance of any slope. For gaze displacement, the covariate reversed the slope’s sign in Monkey J and made it significant in Monkey K, because gaze displacement moved the PSE and the RT asymmetry in opposite directions (Fig. 4C, F).

For hand displacement and head rotation, the slopes were positive and significant in both animals (Fig. 5C–F). In Monkey J the RT asymmetry rose by 1.8% per deg of PSE under hand displacement (95% CI [1.2, 2.4]; p < 0.001; n = 31 shift levels over 11 sessions) and by 1.7% under head rotation (95% CI [1.1, 2.4]; p < 0.001; n = 28 shift levels over 10 sessions). In Monkey K the RT asymmetry rose by 2.4% under hand displacement (95% CI [2.0, 2.9]; p < 0.001; n = 24 shift levels over 8 sessions) and by 1.9% under head rotation (95% CI [1.1, 2.8]; p < 0.001; n = 14 shift levels over 5 sessions).

Without the gaze displacement as a covariate (p values uncorrected), the slope was negative in Monkey J (-5.0% per deg of PSE; 95% CI [-7.2, -2.7]; p < 0.001) and not significant in Monkey K (+1.2%; 95% CI [-0.48, 2.8]; p = 0.16). Within each gaze condition the slope was positive in both animals, from 0.61 to 3.5% per deg in Monkey J and from 2.0 to 4.3% in Monkey K, though in Monkey J only the centered condition was significant. With the gaze displacement entered as a covariate, the slope was +1.3% per deg in Monkey J (95% CI [-0.66, 3.3]; p = 0.20; n = 15 shift levels over 5 sessions) and +2.7% in Monkey K (95% CI [1.9, 3.5]; p < 0.001; n = 21 shift levels over 7 sessions; Fig. 5G, H).

We also tested whether the slope of the RT asymmetry on the PSE changed with the shift, by adding a shift × PSE interaction to the same model (Supplemental Results). After correction for multiple comparisons, the interaction was significant for head rotation in Monkey J, whose slope grew steeper as the head rotated rightward (adjusted p = 0.02). For hand displacement in Monkey J the interaction was also significant (adjusted p = 0.048), though not when RT was summarized by its median (Methods). The interaction was not significant for the other four combinations of manipulation and animal.

## Discussion

In both animals, hand choice depended on horizontal target position. The left hand became more likely to be chosen as targets moved leftward, and the right hand as targets moved rightward. The PSE lay at or just to the right of the body midline. Hand displacement, head rotation, and gaze displacement each moved the PSE in the direction of the manipulation. Reaction times were longest near the PSE, where the hands were chosen about equally often, and the right-versus-left RT asymmetry covaried with the PSE. The sigmoidal dependence on target position, the PSE shifts with head rotation and gaze displacement, and the near-PSE RT increase are also features of human hand choice, so the macaque is a tractable model for it, and the PSE characterized here can now be studied directly at the neural level.

### Reference frames

Target position can be defined relative to the body, the head, the eyes, or the hand, and hand choice could depend on target position in any one of these reference frames or in several. Each manipulation changed target position in one frame and left the others unchanged. Head rotation, with gaze held on the central fixation point, changed target position relative to the head; gaze displacement, with the head still, changed it relative to the eyes; and hand displacement changed it relative to the hand. Hand displacement also changed the distance the hand must travel to each target, so its effect could reflect either target position relative to the hand, as in the hand-centered code proposed for reaching (Graziano 2001), or the biomechanical cost of each reach, as in cost-based models of hand choice (Schweighofer et al. 2015), and our design cannot distinguish the two.

Because head rotation and gaze displacement are angles, as the PSE is, the PSE shift each produced can be compared directly with the rotation or displacement itself. A purely head-centered choice predicts that the PSE moves about as far as the head is rotated and not at all with gaze, a purely eye-centered choice predicts the reverse, and a choice centered on the body or the hand predicts that the PSE moves with neither. Instead, both moved the PSE in the direction of the manipulation, but by only a fraction of the rotation or displacement (head slopes, 0.23 and 0.49 deg per deg in Monkeys J and K; gaze slopes, 0.16 and 0.12). The head slope exceeded the gaze slope in both animals, but the confidence intervals separate the two only in Monkey K, so we treat this ordering as suggestive. Human hand choice also shows partial shifts, with equal head and gaze slopes (Bakker et al. 2018). Neither result fits a single frame, which suggests that hand choice draws on target position in more than one.

If hand choice draws on target position in more than one reference frame, the neurons that select the hand are likely to receive spatial information in several frames at once. The posterior parietal cortex is a strong candidate. In humans, transcranial magnetic stimulation of the posterior parietal cortex biases hand choice (Oliveira et al. 2010), and in macaques, neurons in its parietal reach region code reach targets in eye-centered, hand-centered, or intermediate frames (Chang and Snyder 2010). A choice read out from such a population could shift with both gaze and hand position, each by a fraction, as the PSE did. Recording from these neurons during free choice could test this, and could also separate a hand-centered effect of hand displacement from a biomechanical one, because a hand-centered code would move a neuron’s spatial tuning with the hand whereas a reach cost need not.

### Reaction time and hand choice

Two RT results bear on how the hand is chosen, and they constrain it to different degrees. The first is an increase in RT near the PSE. In human reaching, RT to central targets is longest when the hand is chosen freely and shortest when it is assigned in advance (Oliveira et al. 2010), and a brief train of electrical pulses to one wrist, which shifts choices toward that hand, shortens RT to near-PSE targets (significantly only for the left wrist) while leaving RT to peripheral targets unchanged (Hirayama et al. 2024). Both results tie the increase to choosing the hand. We report a similar increase for reaches near the PSE (Fig. 3). This increase is suggestive of a decision process that takes longer for reaches near the PSE, possibly due to mutual inhibition, but could also be independent of the choice, for example if the animal valued reaches to those targets less, regardless of which hand was chosen.

The second result, the correlation of the RT asymmetry with the PSE (Fig. 5), is stronger evidence that hand choice and reaction time depend on a common process. The asymmetry was measured at peripheral targets, where the hand on the target’s side made 95.1 to 99.8% of reaches in every animal and condition, so those reaches carry almost no information about where the PSE lies (Methods). The correlation therefore cannot come from slow reaches near the PSE, and it shows that RT varied with the PSE even on reaches that almost always used the same hand. With the asymmetry measured only at targets at least 21 deg from the midline, more than 6 deg from every session’s PSE, each slope stayed positive, changed by at most 25%, and kept its significance. The correlation does not show whether the hands inhibit each other or race independently. Under either architecture a hand whose evidence accumulates more slowly both responds later and is chosen less often (Krajbich et al. 2015).

Gaze displacement is the strongest available test of the correlation, because it was the only manipulation that moved the two measures in opposite directions. Displacing gaze rightward moved the PSE rightward but shortened the right hand’s RT relative to the left hand’s (Fig. 4C, F). Across sessions without a manipulation the relation has the opposite sign, with a rightward PSE accompanied by a relatively slower right hand. A correlation produced by the manipulations would have had the sign of the manipulation’s effect. Instead, every per-condition gaze slope was positive (Supplemental Table S1), as for the other manipulations. Fitted across conditions with the shift as a covariate, the correlation was significant in Monkey K (2.7% per deg, p < 0.001) but not in Monkey J (1.3%, p = 0.20). The correlation is therefore unlikely to be an artifact of the manipulations. The RT asymmetry is also not a function of the PSE alone, because then no manipulation could have moved the two in opposite directions.

Why gaze moved the two measures in opposite directions is unresolved. A PSE set partly by gaze moves with the eyes, so a rightward gaze displacement assigns part of the space just right of the midline to the left hand. RT may instead depend on the target’s position relative to fixation, decreasing for targets nearer the point of gaze whichever hand reaches them. Two mechanisms could produce that dependence. Targets near the direction of gaze may be processed faster, by orienting or by attention, so a reach to them starts sooner regardless of which hand is used. The dependence may instead be motor, if transforming a target from retinal to limb coordinates takes longer the farther the target lies from fixation; saccade direction does affect eye-hand coordination in bimanual reaching (Mooshagian et al. 2014). The two accounts differ in whether the shorter RT depends on the target’s position or on the hand used. Our data cannot separate them, because each hand’s RT was measured only at targets on its own side (Methods), where both accounts predict the same result. Separating them requires a design that decouples target position from reaching hand, such as instructing the hand on a subset of trials in each gaze condition.

The PSE-RT correlation has precedent in saccade choices. Monkeys chose between two saccade targets whose reward values changed from block to block, and most trials showed only one of the targets (Milstein and Dorris 2011). Within a block, when a target appeared alone, RT was faster for the target chosen more often and slower for the target chosen less often. When two targets appeared with staggered onsets, a race model built from single-target RT predicted the onset delay at which the two were chosen equally often, the analog of our PSE (Adam et al. 2019). In both saccade studies, RT came from single-target trials, on which there was nothing to choose, and reward or onset delay moved a choice between movement goals rather than hands. In our data the PSE varied across sessions with the reward equal for the two hands, it moved with the position of the hands, head, and eyes, and the slope of the RT asymmetry against it was positive in every condition.

### Relation to prior macaque work

Earlier work on macaque hand choice found that individual animals had a hand preference (Dancause and Schieber 2010; Gardinier et al. 2006; Lee and Schieber 2006), that the target’s side had only a small effect on hand choice (Dancause and Schieber 2010; Gardinier et al. 2006), and that animals more often chose the hand on the side the head faced (Dancause and Schieber 2010). These studies had the animals retrieve food from two fixed wells, one on each side, with free hand choice interleaved among trials that instructed the hand or the target. Two target locations are enough to show that choice depends on the target’s side and that head direction shifts it, but not to measure how choice varies with target position across the workspace, or where the two hands are chosen equally often. We presented targets at ten horizontal positions, never instructing the hand or imposing a delay before the reach, and measured reaction time on the same trials. This design gave the full psychometric function, its PSE, and the reaction-time asymmetry that varied with the PSE. Our head and gaze manipulations also shifted choice in the opposite direction to the earlier head-direction effect. Turning the head or gaze to the right moved the PSE rightward, so the left hand was used for more of the targets near the midline (Fig. 2G–I). The earlier effect was strong in only one of two monkeys, in a task where the choice was made after a delay (Dancause and Schieber 2010).

The same study is also the closest prior test of the PSE-RT correlation (Dancause and Schieber 2010). It averaged reaction, reach, and grasp times from fully instructed trials per session for the four hand and target combinations, and correlated those averages with how often the animal chose each combination on free-choice trials. Execution times correlated with choice frequency. Combinations chosen more often were executed faster, in grasp time for one animal and in reach time for the other. Reaction times did not correlate with choice frequency. That negative result does not conflict with the correlation we report, because the two measurements were computed differently in three ways. Their four combinations include the two crossed reaches (each hand to the target on its opposite side), so biomechanical differences between crossed and uncrossed reaches affect their timing comparison; our asymmetry compares each hand only at targets on its own side of the body midline. Their correlations used absolute times; ours uses a ratio computed within each session or shift level, because absolute reaction time varies across them. And their times came from instructed trials, where the reaction-time effects of hand choice are weaker (Dexheimer et al. 2022; Oliveira et al. 2010).

### Comparison with humans

Hand choice depended on horizontal target position in humans as well, and in both species that dependence was fitted with a sigmoidal function (Bakker et al. 2018; Stoloff et al. 2011). One human study is directly comparable in scale (Bakker et al. 2017). It measured target angle over a 30 cm radius from the midpoint between the hands, with the hands 9 cm either side of the body midline as ours were, and our angles are subtended over a 30 cm viewing distance about that same midline, so a degree spans the same lateral distance in both. Whereas the mean PSE in that study lay left of the midline during whole-body motion (−5.7 deg, SD 5.8, n = 12), our PSEs lay right of it (0.29 and 4.2 deg). Our 10–90% transition widths (11–13 deg) fall within the human range and below its group mean. That study reports a cumulative-Gaussian σ of 7–8 deg (a 10–90% width near 19 deg), with individual widths from roughly 8 to 42 deg.

### Handedness

A preference for one hand moves the PSE toward the other side, because the favored hand is also used for some targets across the midline. Humans are strongly right-handed, a preference that reflects performance differences between the arms (Przy-byla et al. 2013; Sainburg 2002) and appears even in reaction times under free choice (Dexheimer et al. 2022). Consistent with this preference, the PSE of right-handed participants lay on average left of the midline, both with the body stationary (Stoloff et al. 2011) and during passive whole-body translation (Bakker et al. 2017). Reports of hand preference in macaques vary across individuals and across tasks (Chatagny et al. 2013), and several describe a left-hand reaching bias (Fagot et al. 1991; Papademetriou et al. 2005). The PSEs of Monkeys J and K lay right of the midline, the direction that bias implies, with the interval excluding the midline in Monkey K. Monkey A’s PSE also lay to the right. Its start target was never placed left of the midline, but moving it rightward did not move the PSE rightward (Supplemental Figure S2G). The agreement in direction is consistent with those reports but does not by itself establish a macaque population offset, particularly given the individual variation they describe, which our two animals also showed (Monkeys J and K differed in the PSE).

Hand preference had a much larger effect in a food-retrieval task, in which monkeys often used the preferred hand even for targets 30 deg from the midline on the opposite side (Gardinier et al. 2006). The authors suggested that retrieving pellets from deep wells with fine finger movements guided by touch favored the preferred hand; our animals only touched a target on a screen. Humans likewise use the preferred hand more often when a task demands more skill than a simple reach (Bryden et al. 2011), even for targets on the opposite side (Gabbard et al. 2003). Our data do not test whether task demands explain the difference between the studies.

### Translational relevance

Understanding hand choice is relevant to clinical populations in which effector choice is impaired or pathologically biased, such as “learned non-use” of the contralesional hand after unilateral stroke (Kim et al. 2022; Taub et al. 2006). Because shifting the hands, head, or gaze toward one side favored the hand on the other, shifting any of them toward the ipsilesional side could be tested as a way to encourage use of the contralesional hand, rather than constraining the ipsilesional hand.

### Limitations

We tested a limited set of manipulations, one variable at a time; combining hand, head, and gaze within sessions will be needed to determine whether these signals add, interact, or combine according to formal cue-integration principles. The main results also come from two animals, and Monkey K’s head-rotation data from five sessions. The opposing between-condition trends for gaze, in particular, warrant targeted follow-up. Because fixation was enforced only for those 50 ms, the gaze manipulation set where gaze started on each trial, not where it was when the hand was chosen.

## Materials and Methods

All procedures conformed to the *Guide for the Care and Use of Laboratory Animals* and were approved by the Washington University Institutional Animal Care and Use Committee. The analysis code was written with the assistance of the AI tools named in the Acknowledgments, and the authors checked its results.

### Subjects

Two male rhesus macaques (*Macaca mulatta*), Monkey J (16 kg, 10 yr) and Monkey K (9.5 kg, 10–11 yr), participated in the main experiments. A third male animal (Monkey A, 13 kg, 22 yr), naive to home-button and bimanual reaching, was tested in a modified task (Supplemental Figure S2). Each animal was implanted with a head post under aseptic conditions.

### Apparatus

During experiments, animals sat in a custom primate chair with an open front allowing unrestricted reaching with both hands. The head post held the head fixed throughout each session, facing straight ahead or, in head-rotation sessions, turned about the vertical axis to −15 or +15 deg. A 32-inch acoustic touch-sensitive monitor, mounted vertically (in the frontal plane), was positioned 30 cm from the animal’s eyes, with the screen center approximately 7 cm below eye level. Two capacitive home buttons were separated by 18 cm (left and right of midline), each paired with an LED used to cue hand placement. Eye position was monitored with an ISCAN ETL-200 infrared eye tracker (RK-826PCI, 120 Hz) and calibrated at the start of each session.

### Free-choice reaching task

Each trial began with a central fixation point (a white spot subtending approximately 1.1 × 0.8 deg of visual angle) and illumination of both home-button LEDs. The animal initiated the trial by placing each hand on its own home button and fixating the central point. After a fixed 300 ms fixation hold, a reach target appeared, a filled yellow square approximately 3.3 × 2.5 deg in size (width × height; the display stretched the image horizontally). Fixation was required for a further 50 ms, shorter than the fastest saccadic reaction times reported in monkeys (Fischer and Boch 1983), after which eye position was unconstrained, and the animal released one (and only one) home button to reach the target. Touching the target earned a small, fixed water reward, and holding the touch for 250 ms earned a larger reward, which the experimenter increased over the session to maintain the animal’s motivation. Trials were aborted and excluded if the animal broke fixation prematurely, moved either hand before target onset, released both buttons, or released the target before the hold completed. Reach targets were drawn from a fixed 3 × 10 grid: ten horizontal positions at ±3, ±9, ±15, ±21, and ±27 deg from the body midline, crossed with three vertical positions (−18, −4, and +10 deg relative to the fixation point’s vertical position). Positions are angles subtended over the eye-to-screen viewing distance, with horizontal positions referenced to the body midline rather than to fixation. The target grid was fixed on the screen, so target position is unchanged by all three manipulations below. All analyses treat horizontal target position as the independent variable, pooling across the three vertical positions. Trials were run in blocks of up to 300 (ten repetitions of each of the 30 grid positions, in randomized order); blocks with fewer than 150 completed trials were excluded from analysis. Across included blocks, the number of completed trials averaged 283.8 (range 150–300, n = 258 blocks).

### Manipulations

In separate blocks we manipulated one of three variables, each given a signed value (negative leftward, positive rightward) that was held constant within a block and varied across blocks. For the hand manipulation, the home-button pair was displaced laterally as a rigid unit, preserving the 18 cm separation, by ±3.5 or ±4.5 cm relative to the midline (every session used one of the two). We rotated the head in yaw to −15, 0, or +15 deg with the fixation point held at the screen center, and we moved the fixation point to −15, 0, or +15 deg. Only one manipulation type was tested per day, with one exception (Datasets and sample size). Within a session, blocks followed a Latin-square counterbalanced order over the shift conditions (leftward, centered, rightward), with shift magnitude randomized across cycles (subject to the per-block trial-count inclusion criterion). Every hand-displacement session included both a leftward and a rightward shift. Leftward and rightward head and gaze shifts were balanced within each animal’s dataset. Head rotations required a new eye calibration before the subsequent block; hand and gaze displacements did not.

### Design

We used a within-subject, repeated-measures design; each session was a single day’s recording. Monkeys J and K performed the task both with no manipulation and with all three manipulations; Monkey A performed a variant of the free-choice task without home buttons (see Results): between reaches the animal touched a single start target, with six or eight outward targets placed symmetrically about it at eccentricities of 12 to 28 deg. That start target and its array were repositioned laterally between blocks, in no fixed order, over six positions from 0 to 20 deg rightward of the body midline. Target positions are expressed from the body midline throughout, so Monkey A’s PSE is in the same coordinate frame as Monkeys J and K’s. Monkey A underwent none of the three manipulations. Blocks of the same configuration on the same day measure the same thing, so trials are pooled and summarized once per configuration per session: once per session without a manipulation, and once per shift level per session under one (the logistic fit that yields a PSE and transition width, the mean log reaction times that yield an RT asymmetry). Statistical inference is carried out on those summaries rather than on trials. Inference treats the session as the independent unit, with a random intercept per session in every regression and with tests and intervals computed over sessions (Confidence intervals and slope tests). One Monkey K session also tested hand displacements of ±1.5 and ±2.5 cm as acclimation to the full displacement. Those four shift levels were left out of every analysis, so that no session contributed more than three. The transition width is the span of target positions over which the probability of a right-hand reach is between 10% and 90%, a standard, model-independent measure of psychometric spread (see Strasburger 2001). Sample sizes reported as *n* are counts of sessions without a manipulation and of session-by-shift-level summaries under one.

### Datasets and sample size

Monkey J contributed 134 blocks (38502 trials) over 37 sessions, and Monkey K 124 blocks (34876 trials) over 35 sessions. Three variables could be manipulated: hand displacement, head rotation, and gaze displacement. On 11 of Monkey J’s sessions (29.7%) and 16 of Monkey K’s (45.7%), none of the three was moved. The hands rested symmetrically about the body midline, the head faced forward, and the animal fixated straight ahead. On the remaining 26 (70.3%) and 19 (54.3%), one of the three was varied across blocks, except on one Monkey K session that varied two, hand displacement and head rotation.

The no-manipulation dataset comprises every block from the first set of sessions. Blocks recorded with the manipulated variable at zero on a session of the second set are not included. They are unmanipulated, but they also serve as the zero level of their manipulation’s own regression, and counting them in both places would leave the two datasets over-lapping. The hand-displacement and head-rotation datasets share one block, the zero level of that Monkey K session. The no-manipulation dataset totaled 28 blocks (7787 trials) over 11 sessions for Monkey J and 49 blocks (13721 trials) over 16 sessions for Monkey K. The manipulation datasets comprised, for Monkeys J and K respectively, 41 blocks (12020 trials) over 11 sessions and 32 blocks (8807 trials) over 8 sessions for hand displacement; 39 blocks (11269 trials) over 10 sessions and 20 blocks (5637 trials) over 5 sessions for head rotation; and 26 blocks (7426 trials) over 5 sessions and 24 blocks (7011 trials) over 7 sessions for gaze.

Of all 73378 trials, 17 (0.02%) were completed but could not be assigned to a hand: a touch was recorded, but the button state at contact showed both home buttons still held, so no release marked the reaching hand. A further 147 had no recorded touch. Neither kind can carry a left or right attribution, so both are excluded from every analysis and from the per-block trial count the 150-trial block criterion is applied to.

Reaction-time analyses used the 73214 trials that remained. Of those, 278 (0.38%) were excluded because the reaction time fell below 100 ms and 32 (0.04%) because it exceeded 1000 ms, 310 in all (0.42%), leaving 72904 trials. Two animals contributed the manipulation data, with statistical power deriving from within-animal replication across many sessions; every analysis was performed separately for each animal, and three were also pooled across the two: the gaze PSE slope, the RT trends under each manipulation, and the near-PSE RT increase.

Monkey A contributed 1870 trials over 5 sessions, each fitted once with its blocks pooled (Design). Every session’s PSE fell within the tested target range. In each session we tested whether the proportion of right-hand reaches rose the farther right the target was, by a Wald test of the logistic slope against zero, and counted the session when the slope was positive at *p* < 0.001.

### Psychometric fitting

For each no-manipulation session and each shift level of each manipulation session we fit a binomial logistic model of right-hand choice,

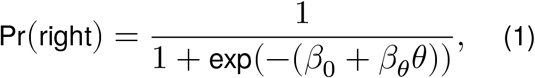

with horizontal target position *θ* (in degrees from the body midline), and defined the PSE as *µ* = −*β*_0_/*β*_*θ*_. The transition width was computed from the slope as *w* = 2 ln(9)/*β*_*θ*_. Extreme negative and positive target positions approach exclusive left- and right-hand use, respectively.

### Reaction time and hand identification

Reaction time was the interval from target onset to the moment the reaching hand first left its home button; the chosen hand was identified from button state at contact: the hand still resting on its home button marked the unchosen hand. Monkey A reached without home buttons, so hand identity could not be read from a button release; instead an experimenter monitored the reaches and set a toggle switch, wired to the same digital inputs, to indicate the hand in use. The toggle was typically set near target contact: relative to target contact, the median setting completed at −7 ms, and 54% of settings completed before contact. We read the toggle state 500 ms after contact, which captures late settings (92% complete by then) while still preceding the return reach on all but 0.05% of trials. Block-level psychometric slope and PSE were indistinguishable across 500–600 ms.

Reaction times shown per target position (Supplemental Figure S9) are geometric-mean reaction times in milliseconds: the exponential of the mean log-transformed RT, taken per target in each session and shift level and then averaged across them. The log transform reduces the right skew of the RT distribution. Reaches averaged 319 ms in Monkey J (interquartile range 285–356 ms) and 246 ms in Monkey K (214–280 ms) across the analyzed conditions.

### PSE-RT correlation

To relate reaction time to hand choice, for each no-manipulation session and each shift level of each manipulation session with a converged psychometric fit we measured the right-versus-left reaction-time asymmetry *A*_RT_,

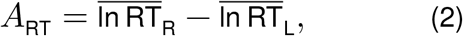

where *R* is the set of right-hand reaches to targets at least 15 deg right of the body midline and *L* the set of left-hand reaches to targets at least 15 deg left of it. *A*_RT_ is the log ratio of their geometric-mean RTs, and it requires at least five trials on each side. The PSE-RT correlation and the trend of each hand’s RT with the manipulation gave the same results when RT was summarized by its median rather than its geometric mean. The correlation also gave the same results when fitted by robust (Huber) regression. With medians, however, the shift × PSE interaction for hand displacement in Monkey J was no longer significant. We report *A*_RT_, and every slope fitted to it, as a percentage, 100 (*e*^*x*^ − 1) for a log-scale quantity *x*, so that 100 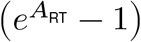 is the amount by which right-hand RT exceeds left- hand RT. We related *A*_RT_ to the fitted PSE *µ* with three linear models, each adding one term to the one before.

Restricting to peripheral targets addresses two biases. First, each trial yields one reaction time, that of the chosen hand. If the decision process is some form of race, then we expect to oversample trials on which the chosen hand was faster (shorter reaction time). This oversampling is smallest at peripheral targets, where the hand on the target’s side is chosen on almost every trial (95.1 to 99.8% in every animal and condition). Any remaining bias would lower the measured asymmetry where the PSE lies to the right and raise it where the PSE lies to the left, flattening the relation, so the slope we report would if anything be an underestimate. Second, the RT asymmetry is measured at peripheral targets, where choices are close to certain and thus have little bearing on the estimate of the PSE. In the no- manipulation sessions, peripheral targets carried a median of only 1.4% (Monkey J) and 5.7% (Monkey K) of each session’s statistical information about the PSE (Fisher information under the fitted logistic function).

#### (i) Per condition

Within a single shift direction (both hand-displacement sizes pooled),

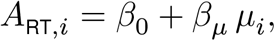

fitted by ordinary least squares to that level’s sessions *i*. These are the per-condition slopes listed in Supplemental Table S1, and they are reported descriptively. Fitted instead to every session and shift level of a manipulation at once, with a random intercept per session as in model (iii), this model mixes the within- and between-condition levels. We report it beside model (iii) for every manipulation (Results; Supplemental Table S2).

#### (ii) Pooled across conditions, adding the shift

Adding the signed shift level *δ*_*i*_ (cm for hand, deg for head and gaze; 0 with no manipulation),

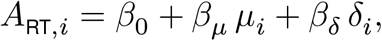

so that *β*_*δ*_ *δ* absorbs the between-condition offset and *β*_*µ*_ is the within-condition slope.

#### (iii) Adding a session intercept

Each session *s* has its own offset *u*_*s*_, modeled as normally distributed across sessions (a random intercept). For shift level *b* of session *s*,

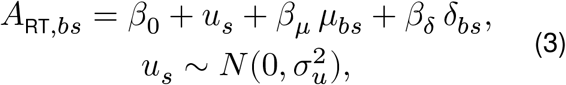

so that the shift levels of one session share an offset rather than entering as independent observations. All three models are reported (Supplemental Tables S1 and S2), and this one gives the PSE-RT correlation quoted in Results under each manipulation: the slope *β*_*µ*_ and its 95% confidence interval as a percentage per degree of PSE, the percentage change in the RT ratio 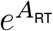 per degree. Without a manipulation each session contributes one point, so there is no session offset to estimate, and the slope is model (i) fitted by ordinary least squares across sessions.

A fourth form, which lets the slope itself change with the shift, is used only for the homogeneity test and is defined with its results (Supplemental Results).

Equation 3 estimates a within-condition relationship. To measure what each manipulation itself did to the RT asymmetry, we regressed *A*_RT_ on *δ* within each session and tested the mean of the persession slopes with the paired sign-flip permutation test described below for each hand’s reaction time. The six slopes, three manipulations in each animal, were corrected together for the false discovery rate (Benjamini–Hochberg procedure, q = 0.05), and the marks in Fig. 4 and the Results give the corrected values.

### Confidence intervals and slope tests

Summaries over the no-manipulation sessions (the median PSE, the median transition width, and the differences between the animals’ medians) are reported as the observed estimate with a 95% bootstrap confidence interval (percentile method, 10,000 resamples of sessions). The confidence bands in Fig. 4 come from 10,000 resamples of sessions. Monkey A’s median PSE is taken over sessions, each pooling blocks that differed in array position. Correlation slopes under the manipulations were estimated by fitting Equation 3 as a linear mixed model. Their confidence intervals and *p* values are the two- sided Wald interval and test for the *β*_*µ*_ term, converted to a percentage per degree (PSE-RT correlation). Where the session variance was estimated at zero, the fit reduced to ordinary least squares. As a stricter check, a model with a fixed intercept per session, which estimates *β*_*µ*_ from within-session variation only, gave a significant slope in five of the six manipulation fits, all but gaze displacement in Monkey J, as the mixed model did (Supplemental Table S2). As a check that does not depend on the model, Supplemental Table S2 also gives cluster-robust intervals, which keep the ordinary least-squares slope but compute its standard error while allowing the residuals of shift levels from the same session to be correlated. With 5 to 11 sessions per fit, that estimator is itself imprecise.

For each manipulation and animal, the PSE at each shift level of each session was regressed on the signed shift with a random intercept per session, as in Equation 3 (fitted by REML, reducing to ordinary least squares where the session variance was estimated at zero), and the slope was tested against zero (no effect of the manipulation on hand choice) by a two-sided Wald test. Head rotation and gaze displacement are measured in degrees of visual angle, the same unit as the PSE, so a slope of one would mean that the PSE moved by the full angle of the manipulation; those two slopes were also tested against a slope of one. Hand displacement is measured in centimeters. Expressing it in degrees at the screen would require an assumption about the choice criterion, so the hand slope is reported and tested in centimeters only. All slopes are reported with 95% confidence intervals. Where one animal’s slope was not significant, we asked whether the two animals’ slopes differed, by adding the shift’s interaction with the animal to the model (one intercept per animal, a random intercept per session), and, where they did not, report the slope pooled across animals with one intercept per animal beside the per-animal slopes. Only Monkey K’s gaze slope fell short of significance, so only the gaze slopes were treated this way.

### Multiple comparisons

Statistical significance was assessed at α = 0.05. The test of each manipulation’s effect on hand choice (one test per manipulation and animal) was pre-specified, and its slope is reported with a 95% confidence interval and without post-hoc correction. The companion test of the head and gaze slopes against a slope of one was not pre-specified. It adds one further test per animal for those two manipulations, four in all, and is likewise reported uncorrected; at the p values obtained, no correction over four tests would change which of them are significant. The PSE-RT correlation was analyzed post hoc, after we observed that the reaction-time asymmetry varied with the PSE in both animals. Its two families of formal tests were each controlled for the false discovery rate (Benjamini–Hochberg procedure, *q* = 0.05; Benjamini and Hochberg 1995): the eight correlation slopes against zero (the Equation 3 slope per manipulation and the no-manipulation slope, each animal) and the six slope-homogeneity tests (Supplemental Results), and Results quote their adjusted p values. The per-condition slopes decompose these correlations by condition and are reported descriptively, uncorrected, with their confidence intervals in Supplemental Table S1.

Each manipulation’s effect on a single hand’s absolute reaction time was measured on the same trials that enter *A*_RT_, taken one hand at a time. Within each session we fitted that hand’s mean log reaction time against the signed manipulation value and, separately, compared its mean log reaction time between shifted and centered conditions; the resulting per-session slopes and differences were then averaged and tested by a paired sign-flip permutation test over all 2^*n*^ sign assignments, so these p values are exact. Pairing within the session holds the block-to-block drift in absolute reaction time fixed, so that drift does not mask the effect. Both tests are reported for every manipulation, and Results give each slope’s size, with its direction stated in words. Because the test has only 2^*n*^ sign assignments over *n* sessions, its smallest attainable two-tailed p is 2/2^*n*^; that floor is quoted wherever a condition’s session count places it above α, so a null result there is read as a limit of the design rather than as evidence of no effect. Each hand’s trend was also tested pooled across the two animals, with the animals compared, as for the near-PSE RT increase (below). These tests were post hoc. The 24 tests for single animals, two per hand for each manipulation and animal, were controlled together for the false discovery rate (Benjamini–Hochberg procedure, q = 0.05). The six pooled trends, two hands under each manipulation, formed one family and their animal comparisons another, each controlled the same way.

The near-PSE RT increase is the percentage by which a hand’s reaction time to targets within 6 deg of the PSE exceeds its reaction time to targets at least 15 deg beyond the PSE on its own side. Four of Monkey K’s points were excluded because the PSE lay more than 12 deg toward that hand’s side, leaving no target 15 deg beyond it. The near-PSE RT increase was tested by the same paired sign-flip test, over each session’s increase without a manipulation and over each session’s mean increase across its shift levels under one; beyond 13 sessions, 10,000 random sign assignments stood in for all 2^*n*^. The primary test pooled the two animals, one test per hand without a manipulation and under each one, over the sessions of both, each session weighted equally. Its 95% confidence intervals are t intervals over the sessions’ mean increases. The animals were compared by reassigning sessions to animals at random. Each of these two families of eight tests was controlled for the false discovery rate (Benjamini–Hochberg procedure, q = 0.05). Each animal was also tested alone, and these 16 tests were controlled together (Supplemental Table S3). We pooled the shift conditions because testing each condition alone would have left Monkey K 4 to 8 sessions per group; pooling matches the no-manipulation tests and asks whether the increase persists under the manipulation.

### Example sessions

Several panels show one session per animal (Figs. 1C, E; 2). These illustrate the fits, and each was selected by a fixed rule. For the no-manipulation panels, the example is the session whose PSE lies nearest that animal’s median. For the manipulation panels, the example is the session with the highest *R*^2^ in the regression of its PSEs on the manipulated variable, among sessions whose slope falls within that animal’s population 95% CI, so that the illustrated session is consistent with the reported group effect. All six manipulation examples satisfied it, each contributing three shift levels.

## Acknowledgments

Claude (Anthropic; Opus, Sonnet, Haiku, and Fable models, versions 4.5 to 5.5) was used to draft and edit the manuscript text, to search and check the literature, and to write analysis code and the code that drew the schematic illustrations. GPT (OpenAI; GPT-5.3-Codex and GPT-5.5) and DeepSeek V4 Pro (DeepSeek) were used to write analysis code. Each was used to speed the work. The authors reviewed and edited all AI-drafted text. The tools were used in a manner that does not conflict with APS ethical policies, and the authors take full responsibility for the content.

## Data Availability

The behavioral data that support the findings of this study, and the custom code used for all analyses, are available from the corresponding author upon reasonable request. All analyses were performed in Python 3.13 (NumPy 2.4, pandas 3.0, SciPy 1.17, statsmodels 0.14, Matplotlib 3.10).

