## Supplementary Material for "Reaction-time asymmetry covaries with shifts in hand choice in macaques"

### Supplemental Results

#### The correlation slope across conditions

For every manipulation, the RT asymmetry rose with the PSE (Results). That slope is fitted within a condition, so a further question is whether the manipulation changes the slope itself. We tested this for each manipulation and animal by adding the displacement, crossed with the PSE, to the correlation model (Methods, model iii),

$$A_{\mathrm{RT},bs} = \beta_0 + u_s + \beta_\mu\, \mu_{bs} + \beta_\delta\, \delta_{bs} + \beta_{\mu\delta}\, \mu_{bs}\, \delta_{bs},$$

where $A_{\mathrm{RT},bs}$ is the RT asymmetry at shift level $b$ of session $s$, $\mu_{bs}$ its PSE, $\delta_{bs}$ its displacement and $u_s$ a session intercept. The test is on $\beta_{\mu\delta}$, by a likelihood-ratio test against the same model without that term. Because $\delta$ enters signed, this tests whether the slope trends with the direction of the shift and has no power against a slope that varies symmetrically about the centered condition.

The statistics were referred to a parametric bootstrap rather than the $\chi^2$ distribution, because the session count (5 to 11) is small. For each manipulation and animal we simulated 10,000 datasets from the maximum-likelihood fit of Equation 3, refit it and the model above to each, and took $p$ as the fraction of simulated statistics at or above the observed one, the observed included. Simulated fits that stopped at a zero session variance were kept. Supplemental Table S1 lists the per-condition slopes and their confidence intervals beside the pooled value they compose.

In Monkey J the slope steepened as the head turned rightward (likelihood ratio $\chi^2(1) = 11.17$, bootstrap $p = 0.004$; Benjamini–Hochberg adjusted $p = 0.02$), as the condition slopes in Supplemental Table S1 show. For the hand manipulation in Monkey J, the interaction was weaker, and in the opposite direction, the slope becoming shallower as the hands moved rightward (likelihood ratio $\chi^2(1) = 7.08$, bootstrap $p = 0.02$; condition slopes +2.1, +2.5 and +1.8% for leftward, centered and rightward), significant after correction for multiple comparisons (adjusted $p = 0.048$), though not with median RTs (Methods).

The remaining tests, Monkey K for all three manipulations and Monkey J for gaze, reached significance on none. A nonsignificant interaction does not show that the slope was constant, so we do not describe any slope as constant. The centered condition of each manipulation is itself an unmanipulated state, recorded in the same sessions as the shifted conditions, so these tests compare manipulated against unmanipulated slopes within a session. We did not compare the pooled slopes against the no-manipulation dataset, which is drawn from separate sessions.

### Supplemental Figures

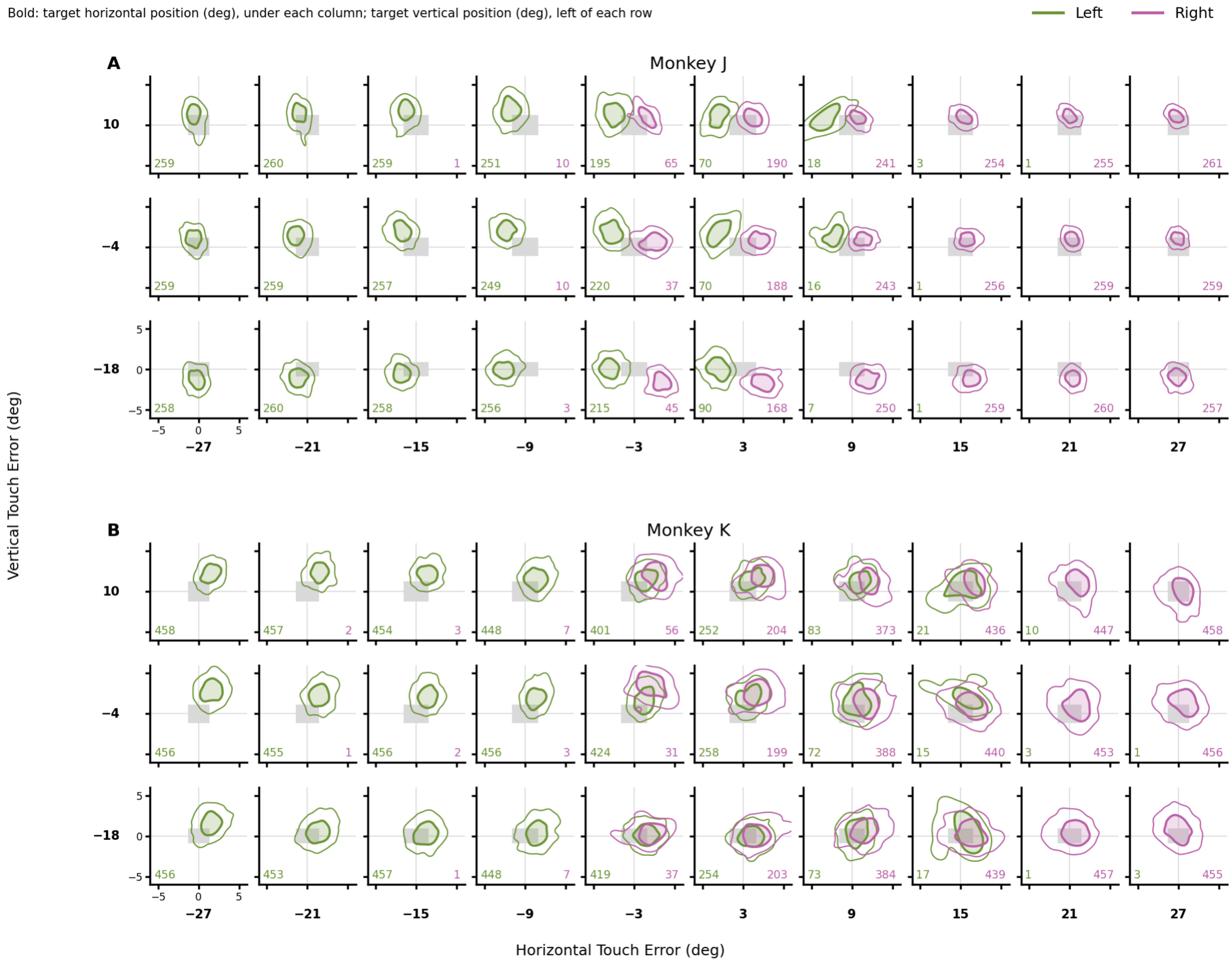


Supplemental Figure S1: **Reaching accuracy, no manipulation.** Where touches landed relative to each target, over all sessions without a manipulation, for Monkey J (A) and Monkey K (B). Each small panel is one target, placed as on the screen and centered on it; gray box, the target, drawn at the angle it subtended at that position. Contours enclose 50% (filled) and 90% of one hand's touches to that target, from a kernel density estimate, drawn where the hand made at least 15 touches; colored numbers, each hand's touch count at that target. In Monkey J, each hand's touches fell toward its own side of the target, the left hand's by up to 3 deg near the midline; in the upper two rows they also fell 1–2 deg above the target, and up to 1 deg below it in the bottom row. In Monkey K, both hands' touches fell 1–2 deg to the right of the target, and 1–2 deg above it in the upper two rows.

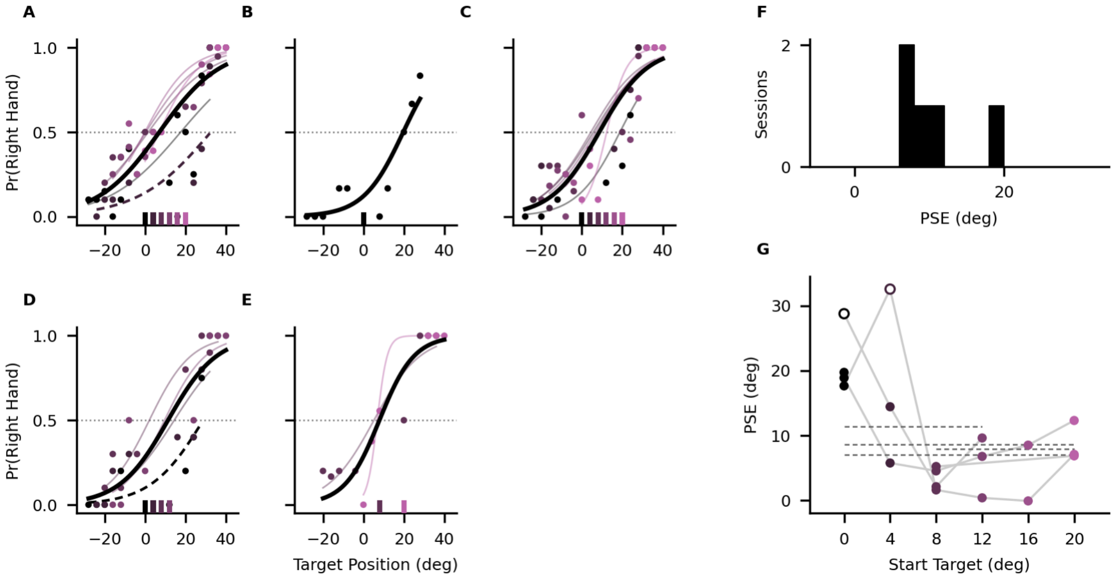


Supplemental Figure S2: **Monkey A, naive to home-button and bimanual training.** (A–E) Proportion of right-hand reaches against horizontal target position, one panel per session in date order. Thick black curve, logistic fit to all of the session's trials, which crosses the dotted line, a proportion of 0.5, at the session's point of subjective equality (PSE). Points, observed proportions, and faint curves, one logistic fit per start-target position, are colored by that position (black, 0 deg, through purple, +20 deg); swatches on the x axis mark the positions the session used. Thin dashed curves, the two start-target fits whose 0.5 crossing lay just beyond the rightmost target tested (A, +4 deg; D, 0 deg). (F) Distribution of session PSEs, each from the thick black curve in A–E (5 sessions). (G) PSE against start-target position, one point per session and position with at least 30 trials. Open points, the two dashed-curve fits of A and D, drawn at their extrapolated crossing. Gray lines join each session's points; dashed lines, that session's PSE from F, drawn across its positions.

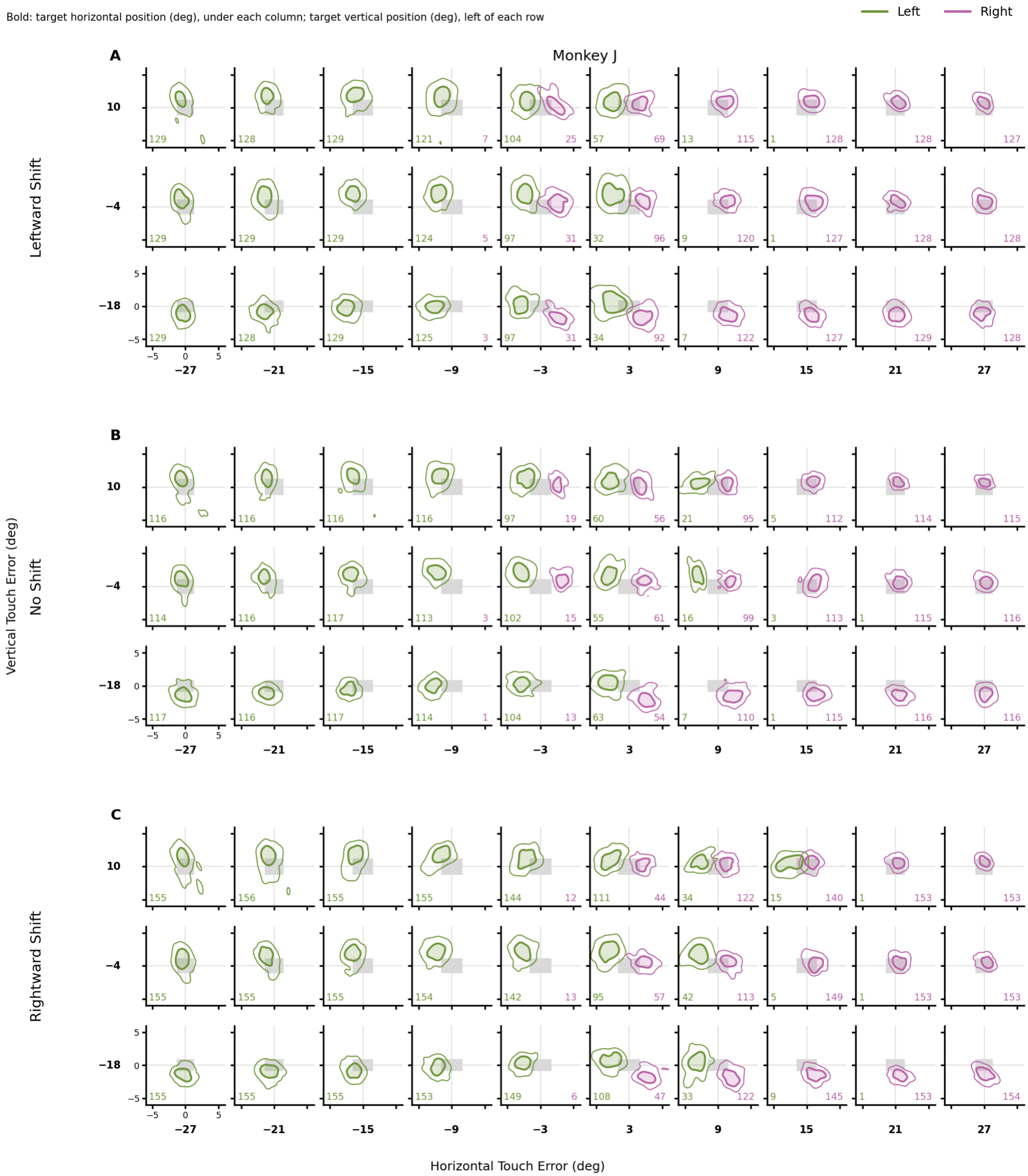


Supplemental Figure S3: **Reaching accuracy under hand displacement, Monkey J.** As in Supplemental Figure S1, pooled over Monkey J's hand-displacement sessions (Fig. 2G), split by the displacement's sign into leftward shift (A), no shift (B), and rightward shift (C). The touch errors follow the pattern described in Supplemental Figure S1 at every shift.

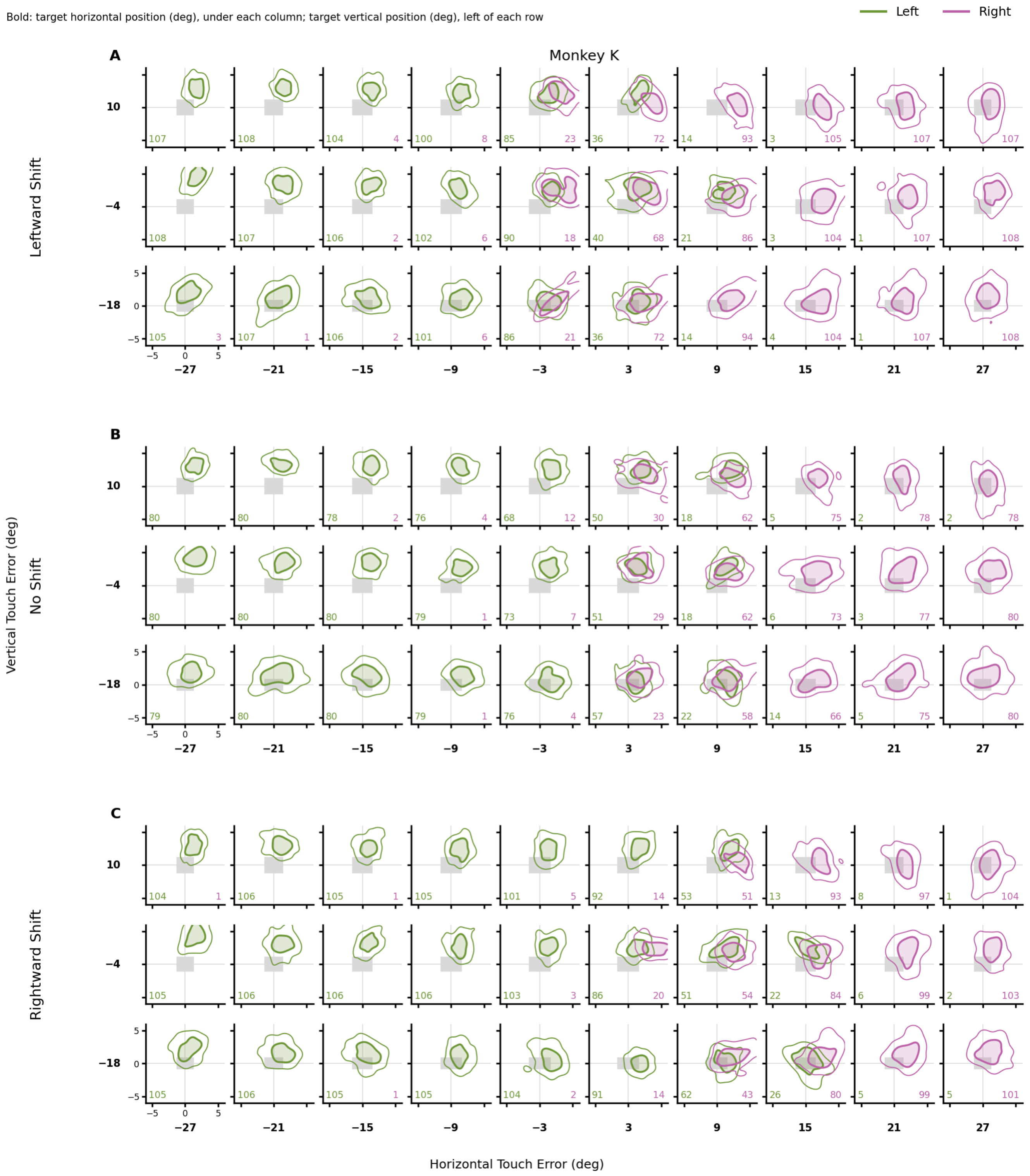


Supplemental Figure S4: **Reaching accuracy under hand displacement, Monkey K.** As in Supplemental Figure S3, for Monkey K. The pattern of touch errors is that of Supplemental Figure S1 at every shift. Monkey K's touches landed about 1 deg higher than in Supplemental Figure S1, with or without a shift.

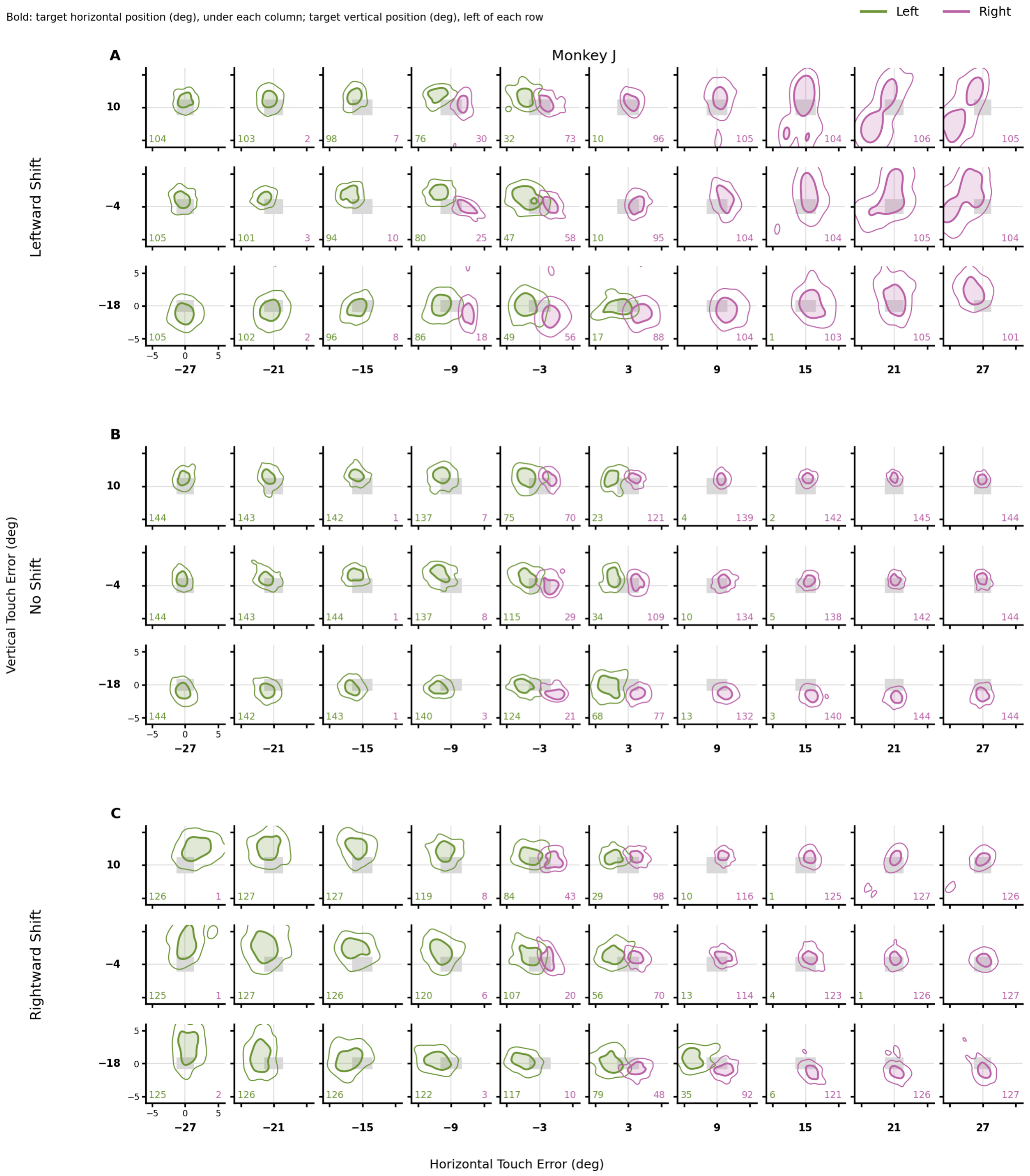


Supplemental Figure S5: **Reaching accuracy under head rotation, Monkey J.** As in Supplemental Figure S3, pooled over Monkey J's head-rotation sessions. With the head rotated, touches to targets on the side opposite the rotation landed higher and scattered more widely than with the head centered (median vertical error, 1.3 vs. 0.7 deg), and about 2% of them fell outside the ±6 deg shown.

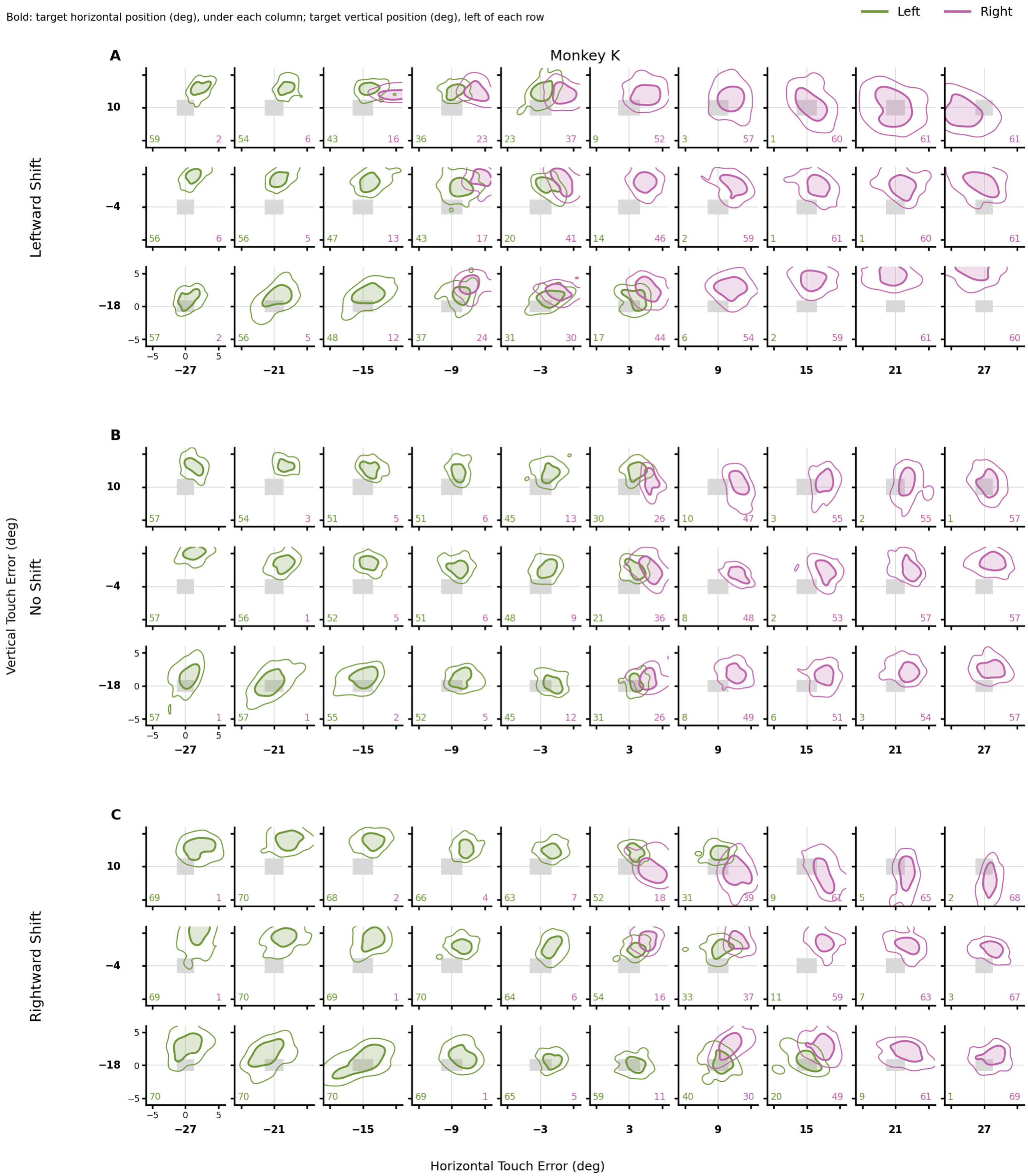


Supplemental Figure S6: **Reaching accuracy under head rotation, Monkey K.** As in Supplemental Figure S3, pooled over Monkey K's head-rotation sessions. With the head rotated, touches to targets on the side opposite the rotation landed higher and scattered more widely than with the head centered (median vertical error, 2.8 vs. 2.1 deg), and about 6% of them fell outside the ±6 deg shown.

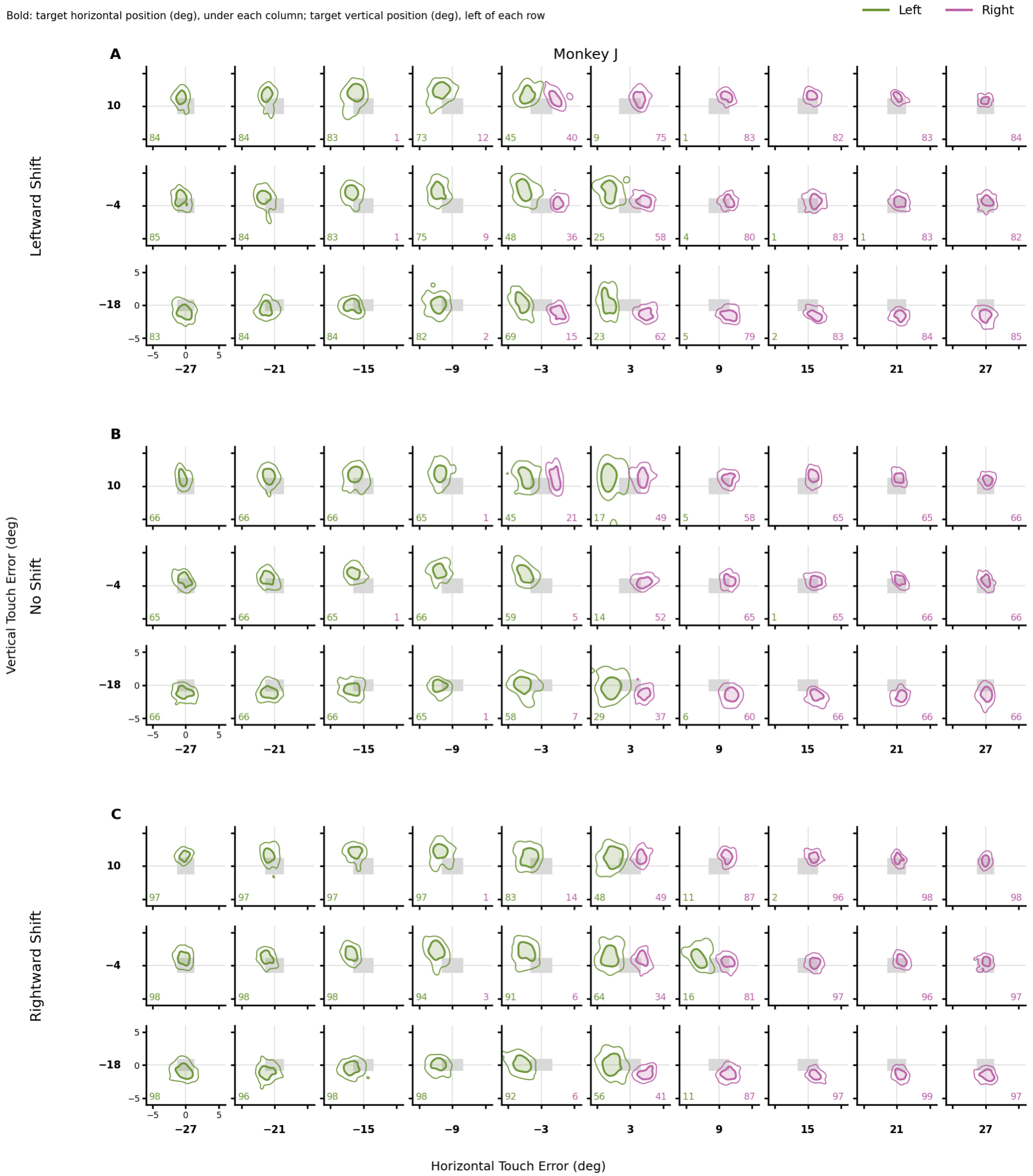


Supplemental Figure S7: **Reaching accuracy under gaze displacement, Monkey J.** As in Supplemental Figure S3, pooled over Monkey J's gaze-displacement sessions. The touch errors follow the pattern described in Supplemental Figure S1 at every shift.

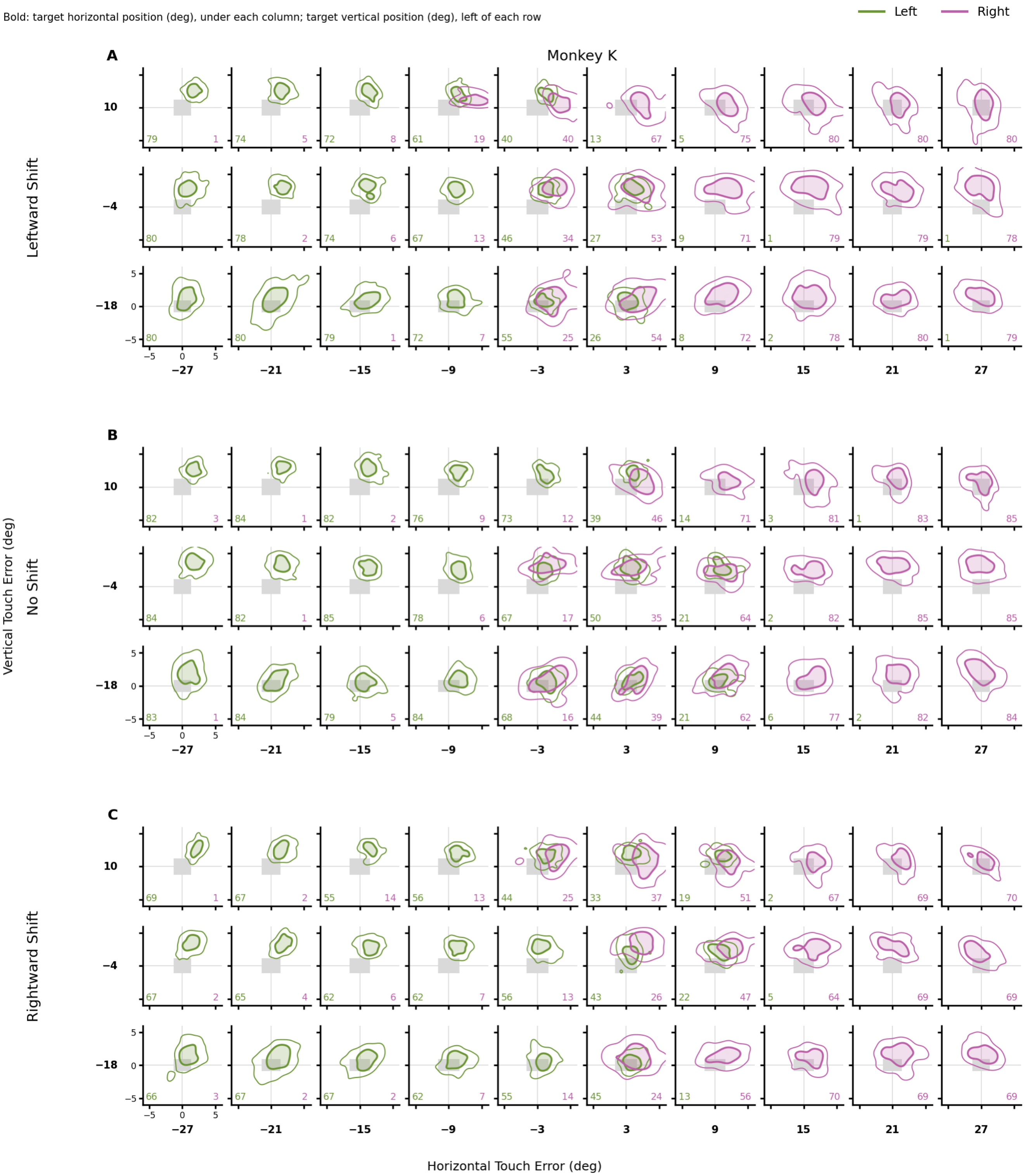


Supplemental Figure S8: **Reaching accuracy under gaze displacement, Monkey K.** As in Supplemental Figure S3, pooled over Monkey K's gaze-displacement sessions. The pattern of touch errors is that of Supplemental Figure S1 at every shift. Monkey K's touches landed about 1 deg higher than in Supplemental Figure S1, with or without a shift.

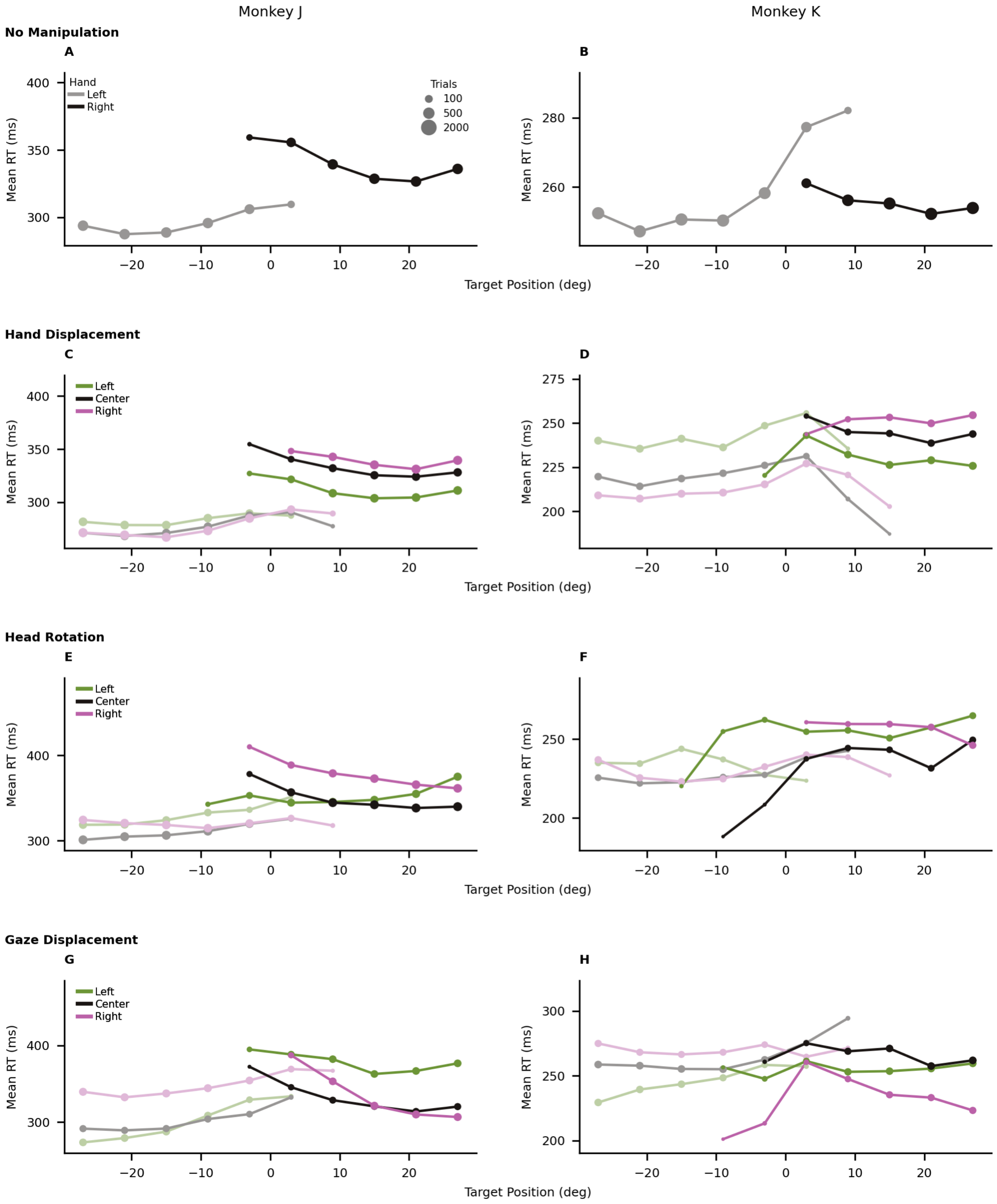


Supplemental Figure S9: **Reaction time by target position.** No manipulation (A, B), hand displacement (C, D), head rotation (E, F), and gaze displacement (G, H), one per row, for Monkey J (left column) and Monkey K (right column). Reaction time (RT), as a geometric mean, against horizontal target position. Pale, left hand; saturated, right hand; green, leftward shift; black, centered; purple, rightward shift. Marker area, trial count (key in A). Points are omitted where a hand made fewer than 10% of reaches. C and D pool the ±3.5 and ±4.5 cm displacements. The sessions are those of Fig. 1D, F and the sessions and shift levels those of Fig. 2G–I.

### Supplemental Tables

Supplemental Table S1: **PSE-RT correlation by condition.** Slope of the RT ratio on the PSE, as a percentage change in the ratio per degree of PSE, for each manipulation and animal. The three condition rows are fitted separately within one shift direction by ordinary least squares; the **All** row is the fit over every session and shift level of that manipulation with the shift as a covariate and a random intercept per session (Methods, model iii) and is the value reported in Results; the partial r is defined only for the per-condition fits. Per-condition *n* is small (fewer than ten for several gaze and head conditions), so those intervals are wide; whether the conditions differ is tested by the shift x PSE interaction reported in Supplemental Results, not by comparing these point estimates. **Same-side % (L / R)** is the share of that row's left-peripheral targets (≥ 15 deg left of the midline) taken by the left hand, and of its right-peripheral targets taken by the right hand; the remainder are the trials the RT asymmetry omits. The two separate where a condition favors one hand, as for head rotation in Monkey K, who used the right hand for 12% of left-peripheral targets when the head was turned left and the left hand for 10% of right-peripheral targets when it was turned right. The lowest values therefore mark the rows where the omission is largest and the slope reported for them most conservative.

| Manipulation | Monkey | Condition | *n* | Same-side % (L / R) | Slope (%/deg) | 95% CI | Partial r | p |
|---|---|---|---|---|---|---|---|---|
| Hand | Monkey J | Left | 11 | 100.0 / 99.8 | 2.12 | [1.25, 3.01] | +0.88 | $p < 0.001$ |
| Hand | Monkey J | Center | 9 | 100.0 / 99.0 | 2.46 | [1.61, 3.32] | +0.93 | $p < 0.001$ |
| Hand | Monkey J | Right | 11 | 100.0 / 97.8 | 1.81 | [0.84, 2.79] | +0.82 | $p = 0.002$ |
| Hand | Monkey J | **All (with covariate)** | 31 | 100.0 / 98.8 | 1.79 | [1.17, 2.42] | — | $p < 0.001$ |
| Hand | Monkey K | Left | 8 | 99.1 / 98.8 | 2.36 | [1.55, 3.18] | +0.95 | $p < 0.001$ |
| Hand | Monkey K | Center | 8 | 99.7 / 95.2 | 2.37 | [1.72, 3.02] | +0.96 | $p < 0.001$ |
| Hand | Monkey K | Right | 8 | 99.8 / 90.6 | 2.47 | [1.04, 3.92] | +0.87 | $p = 0.005$ |
| Hand | Monkey K | **All (with covariate)** | 24 | 99.5 / 94.9 | 2.42 | [1.95, 2.89] | — | $p < 0.001$ |
| Head | Monkey J | Left | 10 | 96.6 / 100.0 | 0.92 | [0.17, 1.68] | +0.71 | $p = 0.02$ |
| Head | Monkey J | Center | 8 | 99.8 / 99.2 | 1.27 | [0.26, 2.28] | +0.78 | $p = 0.02$ |
| Head | Monkey J | Right | 10 | 99.7 / 98.9 | 2.66 | [1.45, 3.88] | +0.87 | $p < 0.001$ |
| Head | Monkey J | **All (with covariate)** | 28 | 98.9 / 99.3 | 1.74 | [1.14, 2.35] | — | $p < 0.001$ |
| Head | Monkey K | Left | 5 | 88.0 / 99.6 | 1.92 | [0.37, 3.50] | +0.92 | $p = 0.03$ |
| Head | Monkey K | Center | 4 | 97.1 / 96.8 | 1.94 | [-2.03, 6.07] | +0.83 | $p = 0.17$ |
| Head | Monkey K | Right | 5 | 99.4 / 89.7 | 1.97 | [0.27, 3.71] | +0.91 | $p = 0.03$ |
| Head | Monkey K | **All (with covariate)** | 14 | 95.0 / 95.1 | 1.90 | [1.05, 2.76] | — | $p < 0.001$ |
| Gaze | Monkey J | Left | 5 | 99.7 / 99.7 | 0.61 | [-4.61, 6.12] | +0.21 | $p = 0.74$ |
| Gaze | Monkey J | Center | 5 | 99.8 / 99.8 | 3.45 | [0.23, 6.78] | +0.89 | $p = 0.04$ |
| Gaze | Monkey J | Right | 5 | 100.0 / 99.8 | 1.12 | [-3.95, 6.46] | +0.37 | $p = 0.54$ |
| Gaze | Monkey J | **All (with covariate)** | 15 | 99.9 / 99.8 | 1.29 | [-0.66, 3.27] | — | $p = 0.20$ |
| Gaze | Monkey K | Left | 7 | 96.8 / 99.4 | 2.99 | [1.18, 4.83] | +0.89 | $p = 0.008$ |
| Gaze | Monkey K | Center | 7 | 98.4 / 98.3 | 4.34 | [1.99, 6.75] | +0.91 | $p = 0.005$ |
| Gaze | Monkey K | Right | 7 | 94.7 / 98.9 | 1.99 | [0.77, 3.22] | +0.88 | $p = 0.008$ |
| Gaze | Monkey K | **All (with covariate)** | 21 | 96.7 / 98.8 | 2.70 | [1.91, 3.50] | — | $p < 0.001$ |

Supplemental Table S2: **The PSE-RT correlation slope without the shift term and under four treatments of session.** Top: percent change in the RT ratio per degree of PSE with 95% CI, for each manipulation and animal. The no-manipulation correlation is not shown: with one PSE per day it has one point per session, so there is no nesting to treat. No shift term is model (iii) with the shift covariate omitted, fitted to all of the condition's shift levels at once, so it mixes the within- and between-condition levels; every other column carries the shift as a covariate (Methods, model ii). OLS treats shift levels as independent. The mixed model adds a random intercept per session and is the fit reported in Results (Methods, model iii). Session fixed effects give every session its own free intercept, so the slope is estimated from within-session variation only. Cluster-robust keeps the OLS estimate and recomputes its interval allowing any correlation among a session's residuals, read against a *t* on sessions minus one. Bottom: p for the shift × PSE interaction (the homogeneity test, Supplemental Results) under an OLS F test, the mixed model's Wald test, the parametric-bootstrap likelihood-ratio test reported in Results (10,000 draws per test), and a cluster-robust *t* test. With 5 to 11 sessions per fit the cluster-robust estimator is imprecise (Methods).

| Condition | Monkey | *n* / sessions | No shift term | OLS | Mixed (reported) | Session fixed effects | Cluster-robust |
|---|---|---|---|---|---|---|---|
| Hand | J | 31 / 11 | 2.37 [1.99, 2.75] | 2.14 [1.65, 2.63] | 1.79 [1.17, 2.42] | 1.34 [0.71, 1.98] | 2.14 [1.54, 2.74] |
| Hand | K | 24 / 8 | 2.72 [2.40, 3.04] | 2.44 [1.98, 2.90] | 2.42 [1.95, 2.89] | 2.36 [1.58, 3.14] | 2.44 [2.04, 2.84] |
| Head | J | 28 / 10 | 1.20 [0.64, 1.76] | 1.74 [1.13, 2.36] | 1.74 [1.14, 2.35] | 2.39 [0.56, 4.26] | 1.74 [1.04, 2.45] |
| Head | K | 14 / 5 | 0.76 [0.12, 1.40] | 1.90 [1.21, 2.59] | 1.90 [1.05, 2.76] | 2.56 [1.31, 3.82] | 1.90 [1.04, 2.77] |
| Gaze | J | 15 / 5 | -4.98 [-7.24, -2.66] | 1.79 [0.03, 3.58] | 1.29 [-0.66, 3.27] | 0.49 [-1.70, 2.74] | 1.79 [-1.38, 5.07] |
| Gaze | K | 21 / 7 | 1.17 [-0.48, 2.84] | 2.66 [1.86, 3.47] | 2.70 [1.91, 3.50] | 2.96 [1.99, 3.95] | 2.66 [1.55, 3.79] |

| Condition | Monkey | OLS F | Mixed Wald | Bootstrap LRT (reported) | Cluster-robust |
|---|---|---|---|---|---|
| Hand | J | p = 0.11 | p = 0.005 | p = 0.02 | p = 0.07 |
| Hand | K | p = 0.26 | p = 0.11 | p = 0.15 | p = 0.12 |
| Head | J | p = 0.002 | p < 0.001 | p = 0.004 | p = 0.08 |
| Head | K | p = 0.43 | p = 0.41 | p = 0.46 | p = 0.49 |
| Gaze | J | p = 0.91 | p = 0.81 | p = 0.83 | p = 0.91 |
| Gaze | K | p = 0.35 | p = 0.32 | p = 0.36 | p = 0.17 |

Supplemental Table S3: **The near-PSE RT increase, per animal and pooled.** For each hand, without a manipulation and under each one, each animal's mean increase over sessions and the two animals pooled, each session weighted equally, with a 95% confidence interval from the t distribution; under a manipulation each session's shift levels are averaged first. p, paired sign-flip permutation test over sessions; with n sessions the smallest attainable p is $2/2^n$. Corrected p, Benjamini–Hochberg over the 16 per-animal tests together and over the eight pooled tests together. Animals differ, sessions reassigned to animals at random, its eight tests corrected together. The pooled test is the one reported in Fig. 3 and Results (Methods).

| Manipulation | Hand | Monkey | Sessions | Near-PSE RT increase, % [95% CI] | p | Corrected p | Animals differ, corrected p |
|---|---|---|---|---|---|---|---|
| No manipulation | Left | J | 11 | 6.9 [4.6, 9.2] | < 0.001 | 0.003 | |
| No manipulation | Left | K | 16 | 9.5 [4.6, 14.7] | < 0.001 | 0.003 | |
| No manipulation | Left | Pooled | 27 | 8.4 [5.5, 11.5] | < 0.001 | < 0.001 | 0.46 |
| No manipulation | Right | J | 11 | 7.6 [4.6, 10.6] | < 0.001 | 0.003 | |
| No manipulation | Right | K | 16 | 3.8 [0.7, 7.0] | 0.004 | 0.008 | |
| No manipulation | Right | Pooled | 27 | 5.3 [3.1, 7.6] | < 0.001 | < 0.001 | 0.18 |
| Hand | Left | J | 11 | 6.1 [3.3, 9.1] | < 0.001 | 0.003 | |
| Hand | Left | K | 8 | 5.5 [2.8, 8.2] | 0.008 | 0.01 | |
| Hand | Left | Pooled | 19 | 5.8 [4.0, 7.7] | < 0.001 | < 0.001 | 0.73 |
| Hand | Right | J | 11 | 3.8 [1.9, 5.7] | < 0.001 | 0.003 | |
| Hand | Right | K | 8 | 2.2 [−0.2, 4.7] | 0.08 | 0.09 | |
| Hand | Right | Pooled | 19 | 3.1 [1.7, 4.6] | < 0.001 | < 0.001 | 0.43 |
| Head | Left | J | 10 | 4.7 [2.2, 7.2] | 0.008 | 0.01 | |
| Head | Left | K | 5 | 3.2 [0.4, 6.2] | 0.06 | 0.08 | |
| Head | Left | Pooled | 15 | 4.2 [2.5, 6.0] | < 0.001 | < 0.001 | 0.46 |
| Head | Right | J | 10 | 3.8 [1.9, 5.7] | 0.002 | 0.005 | |
| Head | Right | K | 5 | 0.9 [−0.5, 2.3] | 0.19 | 0.19 | |
| Head | Right | Pooled | 15 | 2.8 [1.4, 4.2] | < 0.001 | < 0.001 | 0.15 |
| Gaze | Left | J | 5 | 11.8 [7.4, 16.4] | 0.06 | 0.08 | |
| Gaze | Left | K | 7 | 5.4 [1.1, 10.0] | 0.03 | 0.050 | |
| Gaze | Left | Pooled | 12 | 8.0 [4.7, 11.5] | < 0.001 | 0.001 | 0.15 |
| Gaze | Right | J | 5 | 12.4 [5.0, 20.3] | 0.06 | 0.08 | |
| Gaze | Right | K | 7 | 4.5 [−1.2, 10.5] | 0.11 | 0.12 | |
| Gaze | Right | Pooled | 12 | 7.7 [3.3, 12.4] | 0.004 | 0.004 | 0.16 |
